# Listeners with congenital amusia are sensitive to context uncertainty in melodic sequences

**DOI:** 10.1101/2020.07.07.191031

**Authors:** D. R. Quiroga-Martinez, B. Tillmann, E. Brattico, F. Cholvy, L. Fornoni, P. Vuust, A. Caclin

## Abstract

In typical listeners, the perceptual salience of a surprising auditory event depends on the uncertainty of its context. For example, in melodies, pitch deviants are more easily detected and generate larger neural responses when the context is highly predictable than when it is less so. However, it is not known whether amusic listeners with abnormal pitch processing are sensitive to the degree of uncertainty of pitch sequences and, if so, whether they are to a different extent than typical non-musician listeners. To answer this question, we manipulated the uncertainty of short melodies while participants with and without congenital amusia underwent EEG recordings in a passive listening task. Uncertainty was manipulated by presenting melodies with different levels of complexity and familiarity, under the assumption that simpler and more familiar patterns would enhance pitch predictability. We recorded mismatch negativity (MMN) responses to pitch, intensity, timbre, location, and rhythm deviants as a measure of auditory surprise. In both participant groups, we observed reduced MMN amplitudes and longer peak latencies for all sound features with increasing levels of complexity, and putative familiarity effects only for intensity deviants. No significant group-by-complexity or group-by-familiarity interactions were detected. However, in contrast to previous studies, pitch MMN responses in amusics were disrupted in high complexity and unfamiliar melodies. The present results thus indicate that amusics are sensitive to the uncertainty of melodic sequences and that preattentive auditory change detection is greatly spared in this population across sound features and levels of predictability. However, our findings also hint at pitch-specific impairments in this population when uncertainty is high, thus suggesting that pitch processing under high uncertainty conditions requires an intact frontotemporal loop.

## 1. Introduction

Prediction plays an important role in listening. When hearing a melody, for example, we constantly generate expectations about the features of upcoming sounds (Huron, 2006). When contradicted, these expectations give rise to prediction error responses in the brain, which are taken to reflect the update of its internal model of the auditory signal (Friston, 2005; Friston et al., 2020; Vuust et al., 2018). Crucially, the salience of a surprising sound and the strength of the neural responses that it generates are dependent on the uncertainty of the context (Quiroga-Martinez et al., 2019a, 2019b). In the example above, this implies that a wrong or out-of-tune note would be heard more prominently if the melody were repetitive and thus highly predictable. This modulatory effect of uncertainty is often seen as a precision-weighting mechanism that adjusts the balance between top-down predictions and bottom-up sensory signals according to their reliability (Clark, 2013; Feldman & Friston, 2010; Hohwy, 2012).

A growing body of evidence suggests that listeners encode the degree of uncertainty of auditory signals (Barascud et al., 2016; Bianco et al., 2019, 2020; Garrido et al., 2013; Hsu et al., 2015; Lumaca et al., 2019; Quiroga-Martinez et al., 2019a, 2019b; Sohoglu & Chait, 2016; Southwell & Chait, 2018). However, very little is known about how this ability changes in listeners with abnormal auditory function. Studying this type of listeners is of relevance because it could provide valuable information about the auditory processes that underlie the encoding of uncertainty and its modulatory effects. One population of interest in this regard are listeners with congenital amusia, a condition that disrupts the processing of pitch information. Amusic listeners are impaired in pitch discrimination, pitch memory, and pitch production (Ayotte et al., 2002; Graves et al., 2019; Peretz et al., 2002). Given the pitch specificity of this condition, the question arises: do amusic listeners encode the uncertainty of pitch sequences and, if so, do they do it differently than normal listeners?

This question is of particular interest given the neural profile of the condition. Previous research has linked congenital amusia with disrupted functional and structural connections between temporal and frontal areas implicated in auditory sequence processing (Albouy et al., 2013; Hyde et al., 2011; Loui et al., 2009). This is supported by EEG research showing intact early preattentive responses to pitch deviants in amusics, but diminished late responses associated with higher order processing and conscious perception (Moreau et al., 2013; Peretz et al., 2005, 2009; Zendel et al., 2015). These findings have led to the proposal that amusics have normal implicit processing of pitch information, but lack conscious access to it (Norman-Haignere et al., 2016; Peretz, 2016). In this context, a putative abnormal encoding of uncertainty in amusics would suggest that frontotemporal connections play an important role in this function. Conversely, spared uncertainty encoding might suggest its reliance on low-level, preattentive neural processes presumably restricted to the temporal lobe.

However, there is evidence that low-level, implicit processes may also be impaired in congenital amusia. For example, while some behavioral studies have shown that amusics are able to implicitly encode melodic and harmonic expectations when pitch information is task irrelevant (Omigie, Pearce, & Stewart, 2012; Tillmann et al., 2012), they also indicate that the difference between the conditions was smaller than that of controls, which suggests some degree of impairment in implicit processes. Furthermore, research on auditory short-term memory has shown that amusics are impaired, not only in maintenance and retrieval, but also in the encoding of pitch information at the level of auditory cortex (Albouy et al., 2013, 2016, 2019; Tillmann et al., 2016). Specifically, amusics exhibit an abnormal N1 component of the auditory evoked potential (Albouy et al., 2013; Omigie et al., 2013) and show improvements in pitch discrimination when the time to encode pitch information is longer (Albouy et al., 2016; Williamson et al., 2010).

At first glance, this seems at odds with the EEG studies indicating spared low-level pitch processing. However, a potential explanation for the discrepancy is that, in those studies, the stimuli were not complex enough to tax amusics’ sensory memory capacity. For example, Moreau et al. (2013) and Peretz et al. (2005) employed simple, repetitive oddball sequences interrupted by sporadic pitch changes. Similarly, while Peretz et al. (2009) and Zendel et al. (2015) employed several melodies as stimuli, pitch deviants occurred always in the first beat of the third bar, which may have created precise temporal expectations that enhanced deviance salience. This might have made it difficult to detect impairments of implicit processing in amusic listeners. In consequence, manipulating stimulus uncertainty may be crucial to further our understanding of the nature of the deficit.

Another issue where uncertainty manipulations can be particularly informative is whether deficits in amusics are restricted to pitch information, or whether they extend to other musical features. While some studies indicate spared rhythm and beat perception (Graves et al., 2019; Phillips-Silver et al., 2013) and spared use of tempo (Gosselin, Paquette, & Peretz, 2015), timbre (Jiang, Liu, & Wong, 2017), or loudness (Graves et al., 2019) cues, others report impaired contour (Graves et al., 2019) and timbre perception in amusics (Marin, Gingras, & Stewart, 2012; Omigie, Pearce, & Stewart, 2012; Tillmann, Schulze, & Foxton, 2009). Crucially, some evidence suggests that deficits in other features appear once the pitch information—or otherwise spectral information such as timbre—of musical pieces becomes complex. For example, rhythm perception deficits have been reported for variable compared to fixed pitch sequences (Foxton et al., 2006; Pfeuty & Peretz, 2010). Therefore, the question remains whether low-level neurophysiological signatures of the processing of sound features other than pitch are affected by the uncertainty of pitch sequences.

In the present study, we employed EEG to record mismatch negativity (MMN) responses to pitch, intensity, timbre, location, and rhythm deviants from participants with congenital amusia and matched controls, while they listened to melodic contexts with different degrees of uncertainty. The MMN is a well-studied prediction error response to sounds that violate auditory expectations and sensory memory traces (Garrido et al., 2009; Näätänen et al., 1978, 2007; Vuust et al., 2009). We manipulated uncertainty in two ways. First, we changed the complexity of the melodies by varying their repetitiveness and pitch range, as has been previously done (Quiroga-Martinez et al., 2019a). Second, we manipulated familiarity by presenting both well-known children songs and scrambled—but musically plausible—versions of them. These manipulations rest on the assumption that previously known and fairly simple and repetitive tunes provide a source of precise expectations that facilitate deviance detection.

Two alternative hypotheses can be formulated. In one of them, amusics would not be able to encode the uncertainty of the sequences and therefore MMN responses would not change with the complexity or the familiarity of the melodies. In the other scenario, amusics would indeed be able to encode uncertainty, thus inducing a reduction in prediction error responses for more complex melodies and unfamiliar tunes. Crucially, due to the putative higher sensory memory demands of complex and unfamiliar contexts, we conjectured that the effect of increased uncertainty would be stronger in amusics than in controls, resulting in further reductions of MMN amplitudes. Finally, our multi-feature MMN paradigm (Näätänen et al., 2004; Vuust et al., 2011) allowed us to assess whether any observed effects were specific to pitch information, or whether they extended to other auditory features. Thus, if amusics indeed experience preattentive deficits in musical features other than pitch when pitch information becomes complex, a larger uncertainty-driven reduction in MMN responses to intensity, timbre, location and rhythm deviants would be expected for this group compared to controls.

## 2. Methods

The code and materials employed to conduct the experiment and analyses presented here can be found at: https://doi.org/10.17605/OSF.IO/JSEU8.

### 2.1. Participants

Seventeen amusics and 17 matched controls took part in the experiment (see Table 1 for demographics). One amusic participant and their matched control were not included in the familiarity analyses, because the corresponding recorded EEG files were corrupted. All participants were French speakers, were recruited in the Lyon area in France, and were screened for congenital amusia with the Montreal Battery of Evaluation of Amusia (MBEA) (Peretz et al., 2003). The total MBEA scores (6 subtests), *t*(25.6) = −10.486, *p* < .001; and the average scores for the three pitch subtests *t*(23.7) = −11.36, *p* < .001, were significantly lower for amusics than controls (see table suppl. 1 for mean scores of the six subtests). A participant was considered amusic if their total score (i.e. mean number of correct responses) was lower than 23 (maximum score = 30, 77% correct) or their mean score for the pitch subtests was lower than 21.7 (maximum score = 30, 72% correct). Moreover, pitch discrimination thresholds (measured in semitones with an adaptive tracking staircase procedure described in Tillmann et al. 2009) were larger for amusics than controls, *t*(16.4) = 3.375, *p* = .004. No significant differences were found for age *t*(31.9) = 0.145, p = .886, years of education *t*(31.4) = −0.07, p = .946, or musical training U = 127.5, p = .163. The study was approved by a national ethics committee and was conducted in compliance with the Helsinki declaration. Participants gave their written informed consent and received a small monetary compensation.

**Table 1.** Demographic information (mean ± SD). MBEA: Montreal Battery of Evaluation of Amusia; PDT: pitch discrimination threshold. See section 2.1 for corresponding statistical tests.

|  | amusics | controls |
| --- | --- | --- |
| Sample size | 17 | 17 |
| Female | 8 | 9 |
| Right-handed | 13 | 14 |
| Education (years) | 15.06 (± 2.7) | 15.12 (± 2.34) |
| Music training (years) | 0 | 0.24 (± 0.66) |
| Age (years) | 38.43 (± 15.96) | 37.61 (± 17.02) |
| MBEA | 21.79 (± 1.81) | 27.09 (± 1.04) |
| MBEA pitch | 20.79 (± 2.15) | 27.43 (± 1.09) |
| PDT (semitones) | 1.57 (± 1.53) | 0.31 (± 0.17) |

### 2.2. Stimuli

#### 2.2.1. Complexity

We employed pitch sequences with different degrees of melodic complexity (Fig. 1a). All sequences included the same number of standard and deviant tones (see below). Low-complexity (LC) melodies correspond to a modified version of the so-called optimal MMN paradigm (Näätänen et al., 2004), consisting of a repeating piano tone with a fundamental frequency at C3 (≈ 262 Hz). The intermediate complexity (IC) stimuli consisted of a repeated four-note pattern that has been used previously and is referred to as the “Alberti bass” (Vuust et al., 2011, 2012, 2016)^1^. The high-complexity (HC) stimuli consisted of major-mode and minor-mode versions of six non-repetitive melodies (see Appendix 1 for full stimulus set)^2^. See Quiroga-martinez et al. (2019a, 2019b) for details about the stimuli and how complexity—or entropy—was estimated. HC melodies were 32-notes long and were presented eight times in a pseudorandom order, so that no melody was repeated before a whole new instance of the pool of twelve melodies had been played. During stimulus presentation, HC melodies were pseudorandomly transposed from 0 to 5 semitones upwards starting in the key of F major. IC stimuli were transposed in the same way, every 32 notes. In this case, however, melodies were transposed to two different octaves to properly cover a similar pitch range as HC stimuli. LC stimuli were never transposed to minimize uncertainty. Each condition was presented in a separate block lasting approximately 13 minutes. The complexity blocks, together with an additional block involving an MMN paradigm with unpitched sounds (data not presented here), were counterbalanced across participants and their order matched across groups.

**Figure 1.**
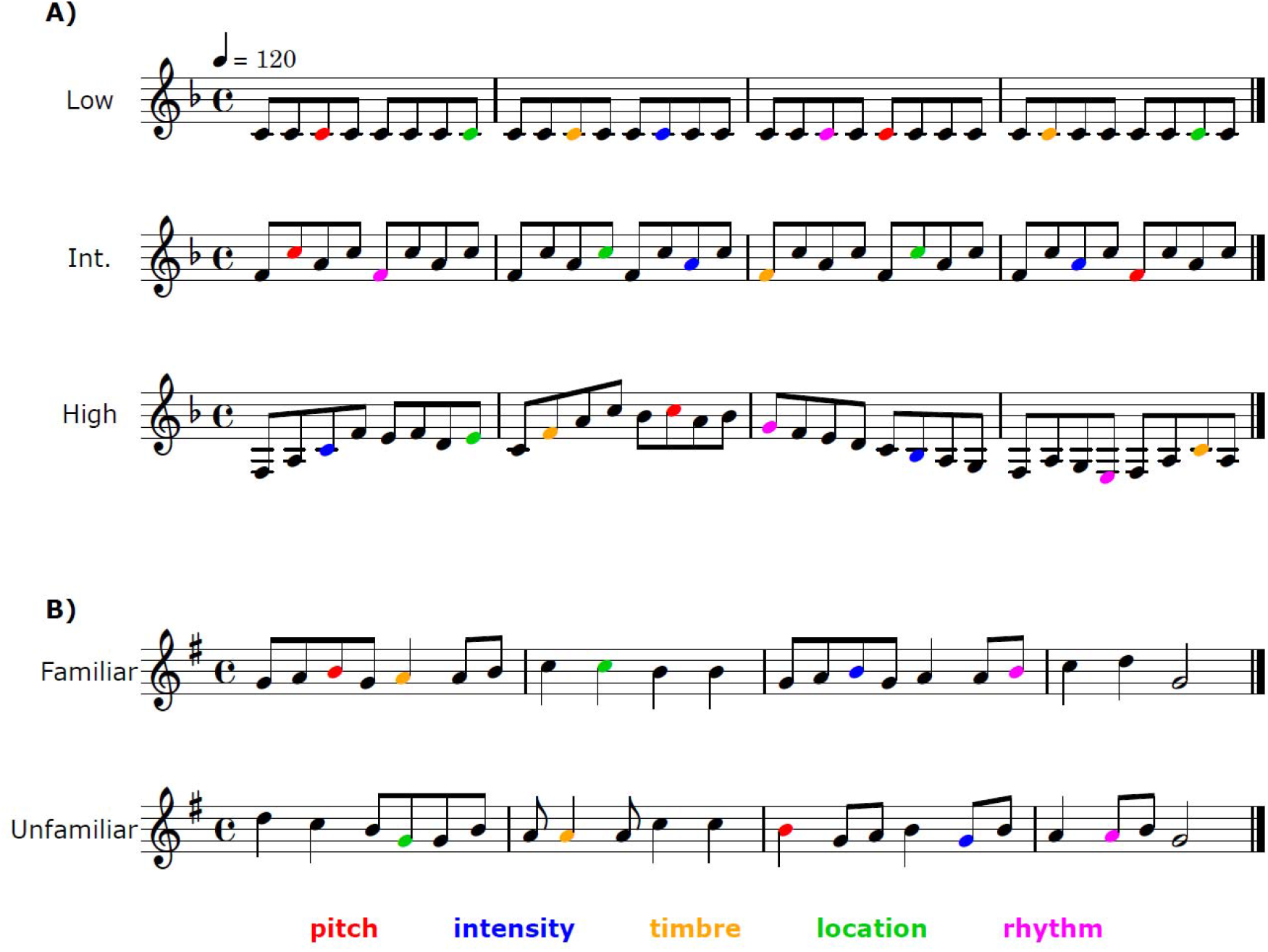
Examples of the stimuli used to manipulate complexity (A) and familiarity (B). Colored notes represent deviants. In the experiment, complexity stimuli were transposed to different keys, whereas familiarity stimuli were always presented in the same key.

The melodies were played with piano tones created in Cubase (Steinberg Media Technology, version 8) with a grand piano sample. The tones lasted 250 ms each and were peak-amplitude normalized. The pitch range spanned 31 different tones, from E3 (F_0_ ≈ 164 Hz) to B□5 (F_0_ ≈ 932 Hz). There were no silent gaps between sounds. Pitch, intensity, timbre, location, and rhythm deviants were introduced in the melodies. These deviants were created with Audition (Adobe Systems Incorporated, version 8) by modifying the standard tones as follows. Pitch: +50 cents; intensity: −12 dB; timbre: a custom notch filter; location: leftward bias (20 ms interaural time difference); rhythm: −60 ms for sound onset. Note that rhythm violations implied a shortening of the preceding tone and a lengthening of the actual deviant tone by 60 ms. Moreover, the magnitude of pitch deviants was chosen to maximize the out-of-tune violation based on previous findings (Quiroga-Martinez et al. 2019a) suggesting that deviants are harder to detect in higher-uncertainty melodies. We incorporated a deviant every four notes in the melodies, with the exact position of the deviant within a four-note group being random. No two deviants were played consecutively and no deviant feature was presented again before a whole iteration of the five features was played. The order of deviant features was pseudorandom. At the start of each block, a randomly selected melody with no deviants was played to properly establish auditory regularities at the outset. A total of 2339 standards and 153 (≈ 5%) deviants per feature were presented for each condition.

#### 2.2.2. Familiarity

Familiar stimuli consisted of seven melodic excerpts from a selection of children’s songs that are well known in France (Fig. 1b) and have been used in previous research (Devergie et al., 2010; Graves et al., 2019). Each excerpt had a 4/4 meter, spanned four bars, lasted 8 seconds and was played in a G major key. Unfamiliar excerpts were created by scrambling the pitch sequence for each familiar tune, with the constraint that they kept the same meter and were musically plausible. Thus, familiar and unfamiliar stimuli were very well matched in their content, only differing in whether the auditory sequence was known by the listener beforehand (see Appendix 1 for the full stimulus set). In a pilot test with 7 French participants, familiar melodies were consistently rated higher than unfamiliar ones on a familiarity scale from 1-7 (Appendix 2).

The melodies were presented using the same type of piano tones as in the complexity conditions, but with different tone durations. The predominant duration was 250 ms, corresponding to eighth notes at 120 bpm. Tones with a duration of 500, 750 and 1000 ms were also included, corresponding to quarter notes, dotted quarter notes and half notes, respectively. A few sixteenth notes with a duration of 125 ms were also present, but these were excluded from the analyses due to their very short duration. The pitch range spanned an octave, from D4 (F_0_ ≈ 294 Hz) to D5 (F_0_ ≈ 587 Hz). There were no silent gaps between sounds. Melodies were pseudo-randomly presented one after the other without pause between them. The order of presentation, the transpositions, the deviants included and their randomization were the same as indicated above. The pool of melodies was repeated 19 times. A total of 2466 standards and 154 (≈ 6%) deviants per feature were included in each block. Familiar and unfamiliar melodies were played in different blocks, with a counterbalanced order across participants that was matched between the groups. Each block lasted around 20 minutes.

### 2.3. Procedure

Upon arrival, participants filled out the required forms, were informed about the procedures and gave their written consent. Then, Biosemi EEG caps with active electrodes were placed on their scalp and conductive gel was applied. A Sennheiser HD280 Pro headset was carefully placed on top of the EEG cap with foam padding to avoid pressure on electrodes. The impedances were checked again after the headphones were on. Sound loudness was set to a comfortable level and was the same for all subjects. During stimulation, participants were sitting on an armchair inside a sound-attenuated booth, electrically-shielded with a Faraday cage, looking at a computer screen from a distance of about 1.5 m. They were informed that there would be sounds playing in the background but were instructed to watch a movie of their choice and ignore the sounds. They were also instructed to remain still and relaxed, but were informed there would be pauses between blocks when they could stretch and change posture. During stimulation, the blocks were presented in such a way that, for 9 matched pairs of participants, the counterbalanced complexity conditions came before the counterbalanced familiarity conditions, whereas for the remaining pairs the order was inverted. Two additional blocks were included at the end of the experiment, in which participants listened freely to entire pieces of music. Their analysis, however, is beyond the scope of this article. The whole recording session lasted around one hour and a half, plus half an hour of preparation.

### 2.4. EEG recording and preprocessing

Scalp potentials were recorded with a 64-channel Biosemi system with active electrodes and a sampling rate of 1024 Hz. Additional electrodes were used to track horizontal and vertical eye movements. Data analyses were conducted with MNE-Python (Gramfort et al., 2014). EEG signals were first cleaned from eyeblink artifacts using independent component analysis with a semiautomatic routine (fastICA algorithm). Visual inspection was used as a quality check. After removing ICA components, the raw signals were filtered with a pass band of 0.5-35 Hz and re-referenced to the mastoids. Epochs −100 to 400 ms from tone onsets were extracted and baseline corrected with a prestimulus baseline of 100 ms. Epochs with an amplitude exceeding 150 μV were rejected to further clean the data from remaining artefacts. For each participant, event-related potentials (ERP) were obtained by averaging epochs for the standard tones and each of the deviant features separately, per condition. Standard tones preceded by a deviant were excluded from the averages. Deviant-specific MMN responses were calculated by subtracting standard from deviant ERPs for each feature and condition.

### 2.5. Statistical analyses

Our analyses comprised two steps. First, we used cluster-based permutations (Maris & Oostenveld, 2007) in a mass-univariate approach, to test differences between standards and deviants, and main and simple effects of complexity and familiarity. This non-parametric method allows for hypothesis testing at the whole-scalp level while controlling the error rate associated with multiple tests. This is achieved by means of spatiotemporal clustering of sample-level statistics, in this case by addition of contiguous samples below a significance threshold of *p* = 0.05. The cluster-level significance of the effects was assessed through Montecarlo sampling of a null distribution obtained from 10.000 random permutations of the labels of the conditions being compared in the test. The resulting *p*-value corresponds to the proportion of sampled cluster-level statistics that were equally or more extreme than the originally observed cluster statistics.

The presence of an MMN for each feature, condition and group was evaluated by comparing standard and deviant ERPs with a two-sided paired-samples *t*-test. The main effect of complexity was assessed for each feature, for amusics and controls combined, by performing an *F*-test on MMN difference waves with the three conditions (LC, IC, HC) as the levels of a single factor. Post-hoc, simple effects of complexity were also tested for each group separately. The main effect of familiarity was assessed for each feature, for amusics and controls combined, by performing two-sided paired-samples *t*-tests between familiar and unfamiliar melodies. Post-hoc, simple effects of familiarity were also tested for each group separately. Given that we tested the same hypothesis for each feature separately, we applied a Bonferroni correction for the main and simple effects of complexity and familiarity by lowering our significance threshold to *p* = .0125 (i.e. 0.05 / 4). Note that Rhythm deviants are excluded from these analyses due to baseline contamination (see results section and Appendix 3 for further details).

In addition to the whole-scalp analyses, in a second stage, we performed analyses on mean amplitudes and latencies. Given that the MMN difference waves peaked at different latencies for different features and conditions, these analyses allowed us to test for feature-specific effects and properly assess main effects and interactions. MMN peak latencies were estimated within a time window of 70-300 ms for pitch, and 70-250 ms for the other features.

Mean amplitudes were obtained from electrodes Fz, F1, F2, F3, F4, FCz, FC1, FC2, FC3, FC4 and calculated as the average activity ± 25 ms around the participant-wise peak, for each participant and feature. The chosen electrodes correspond to those that typically show the largest responses in MMN studies, and exhibited the largest P50 amplitudes (thus making sure that they properly captured auditory evoked activity). Using R (R Core Team, 2019) and the *lme4* library (Bates et al., 2015), several mixed-effects models of mean amplitudes and latencies were compared through likelihood ratio tests, for complexity and familiarity conditions separately. These models were built incrementally from intercept-only models by adding one factor at the time until reaching a full model that included the relevant main factors and their two- and three-way interactions (see table 2). Subject-wise random intercepts were included in all models. No random slopes were introduced, as there were not enough data points to avoid overfitting. Models were also assessed with Akaike Information Criteria (AIC). Post-hoc, pairwise contrasts were also performed with the *emmeans* library (Lenth et al., 2019).

**Table 2.** Likelihood ratio tests for mixed-effects models of mean amplitudes and peak latencies for the complexity conditions. Models were built incrementally by adding a term at the time. Interaction terms are marked with a colon. Comparisons were made between adjacent models (i.e. model vs null). Akaike information criteria (AIC), chi-square statistics (X^2^) and p-values (*p*) are reported. Comparisons with significant differences are highlighted in bold and marked with an asterisk.

| model | added term | null | Mean amplitudes |  |  | Peak latencies |  |  |
| --- | --- | --- | --- | --- | --- | --- | --- | --- |
| | | | AIC | $\chi^2$ | $p$ | AIC | $\chi^2$ | $p$ |
| m0 | Intercept | NA | 1603.9 | NA | NA | 4298.18 | NA | NA |
| m1 | complexity | m0 | 1464.9 | 143 | **<.001\*** | 4269.57 | 32.61 | **<.001\*** |
| m2 | group | m1 | 1466.13 | 0.76 | 0.38 | 4269.96 | 1.61 | 0.2 |
| m3 | feature | m2 | 1419.94 | 52.19 | **<.001\*** | 4049.26 | 226.71 | **<.001\*** |
| m4 | feature:complexity | m3 | 1402.38 | 29.56 | **<.001\*** | 3988.19 | 73.07 | **<.001\*** |
| m5 | group:complexity | m4 | 1403.49 | 2.9 | 0.23 | 3991.02 | 1.16 | 0.56 |
| m6 | feature:group | m5 | 1405.85 | 3.64 | 0.3 | 3988.43 | 8.59 | **0.04\*** |
| m7 | feature:complexity:group | m6 | 1414.45 | 3.4 | 0.76 | 3998.92 | 1.51 | 0.96 |

## 3. Results

### 3.1. Presence of the MMN

Consistent with the literature (Näätänen et al., 2007), the MMN manifested itself as a fronto-central negativity (Fig. 2 and 3). We found differences between standard and deviants for both groups and all conditions for intensity, timbre and location (Fig. 2 and 3). For pitch, the MMN was found in both groups for LC, IC and familiar melodies, but only in controls for HC and unfamiliar melodies. Rhythm MMNs were only detected for IC stimuli in amusics and controls (see Appendix 3 for details).

**Figure 2.**
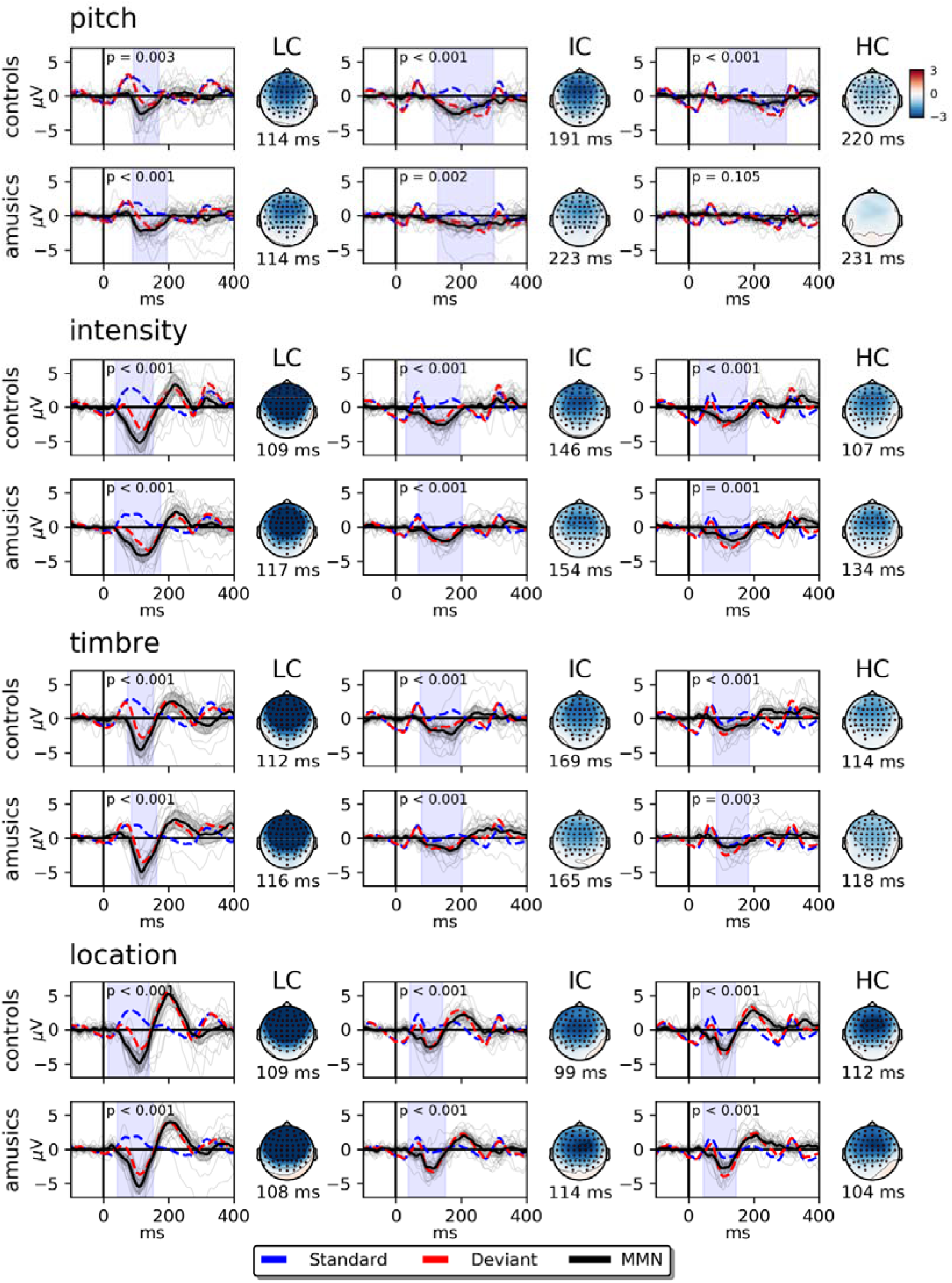
Standard, deviant and MMN difference waves, as well as MMN topographies for the complexity conditions. Activity displayed corresponds to the average of electrodes Fz, F1, F2, F3, F4, FCz, FC1, FC2, FC3 and FC4. Shaded blue areas and red points in the scalp maps mark the times and channels where differences between standard and deviants were significant, as indicated by *p-* values. Shaded grey areas depict 95% confidence intervals. Grey traces depict MMN waves for single participants. LC, IC, HC = low, intermediate and high complexity.

**Figure 3.**
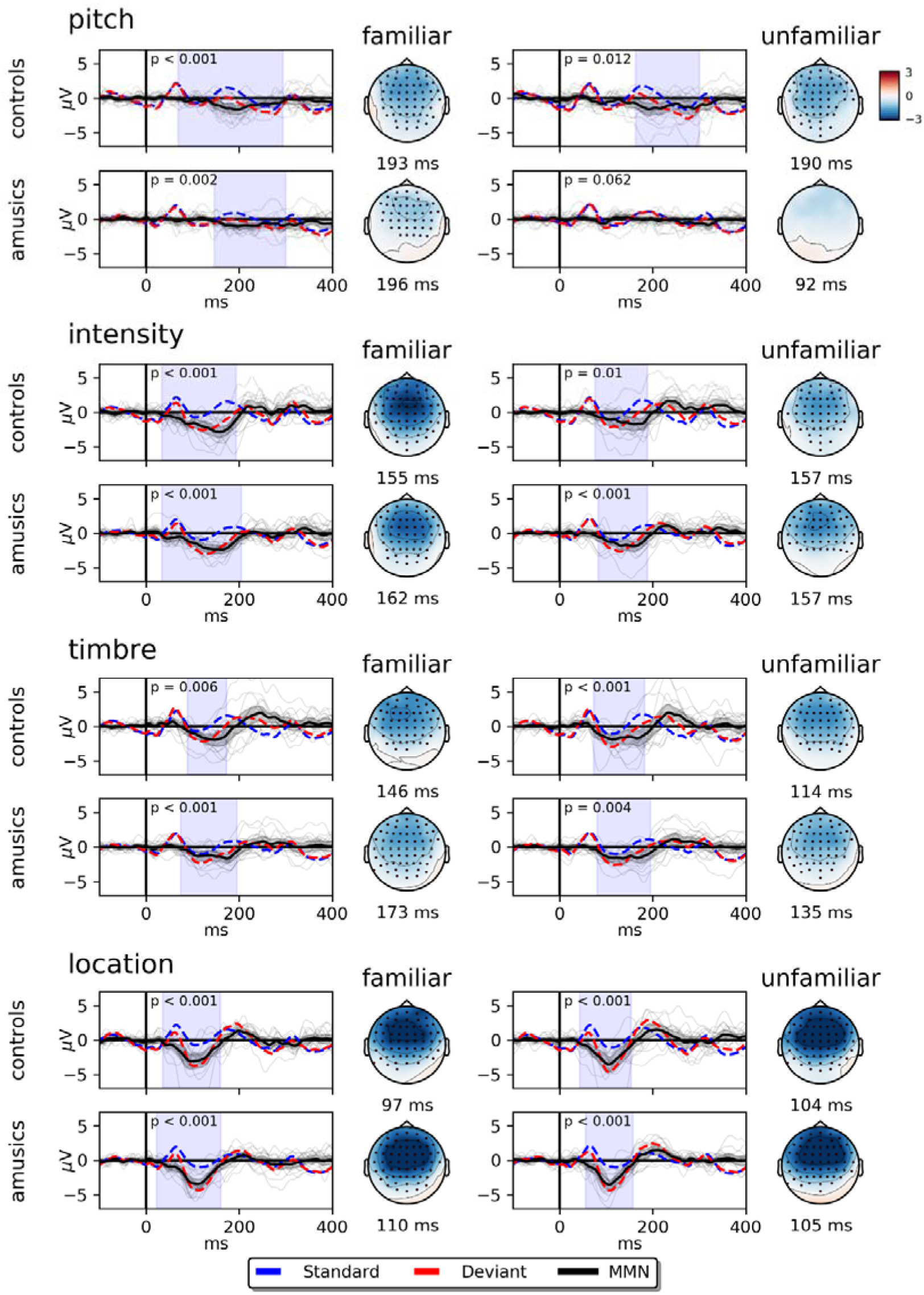
Standard, deviant and MMN difference waves, as well as MMN topographies, for the familiarity conditions. Activity displayed corresponds to the average of electrodes Fz, F1, F2, F3, F4, FCz, FC1, FC2, FC3 and FC4. Shaded blue areas and red points in the scalp maps mark the times and channels where differences between standard and deviants were significant, as indicated by *p*-values. Shaded grey areas depict 95% confidence intervals. Grey traces depict MMN waves for single participants.

In general, the difference waves for rhythm exhibited an abnormal pattern with large peaks before the onset of the current tone, and after the onset of the next tone (Appendix_3). The reason for this is a baseline contamination arising from the different onset times of standard and deviants with respect to the preceding tone in these specific paradigms. Note that the dominant inter-onset interval was relatively short (250 ms), which meant that evoked activity from the preceding tone was still ongoing at target onset. This means that standards and rhythm deviants systematically occurred at different phases of the ongoing activity. This may have affected responses in two ways. First, having a different baseline at the start could have shifted the activity of the deviants with respect to the standards. Second, ongoing activity from the previous tone may have overlapped with the activity of the deviant tone in a way that differed from standard tones. Here we tried to minimize the first by computing and subtracting a specific baseline (−160 ms to −60 ms) from the standard tones to be compared with the rhythm deviants. Despite these corrections, rhythm MMNs were hardly detectable. For this reason, we excluded rhythm deviants from further analyses (Appendix 3).

### 3.2. Complexity effects

Mass univariate analyses revealed significant main and simple effects of complexity on MMN amplitudes for all features for both groups (Fig. 4 and see Appendix 4 for a display of main effects across groups). For intensity, timbre, and location, two clusters were identified, one between approximately 50 ms and 150 ms (corresponding to the MMN latency range), and another one between 150 and 250 ms (corresponding to the P3a latency range). The effect of complexity was confirmed in mixed-effects modeling of mean amplitudes, which revealed that adding the three complexity conditions (*m*1) significantly improved model performance (Table 2; Fig. 4). Interestingly, adding terms for feature (*m*3) and a feature-by-complexity (*m*4) interaction also improved model performance. There was no evidence for an effect of group or group-related interactions. AIC values indicate *m*4 as the winning model (Table 2).

**Figure 4.**
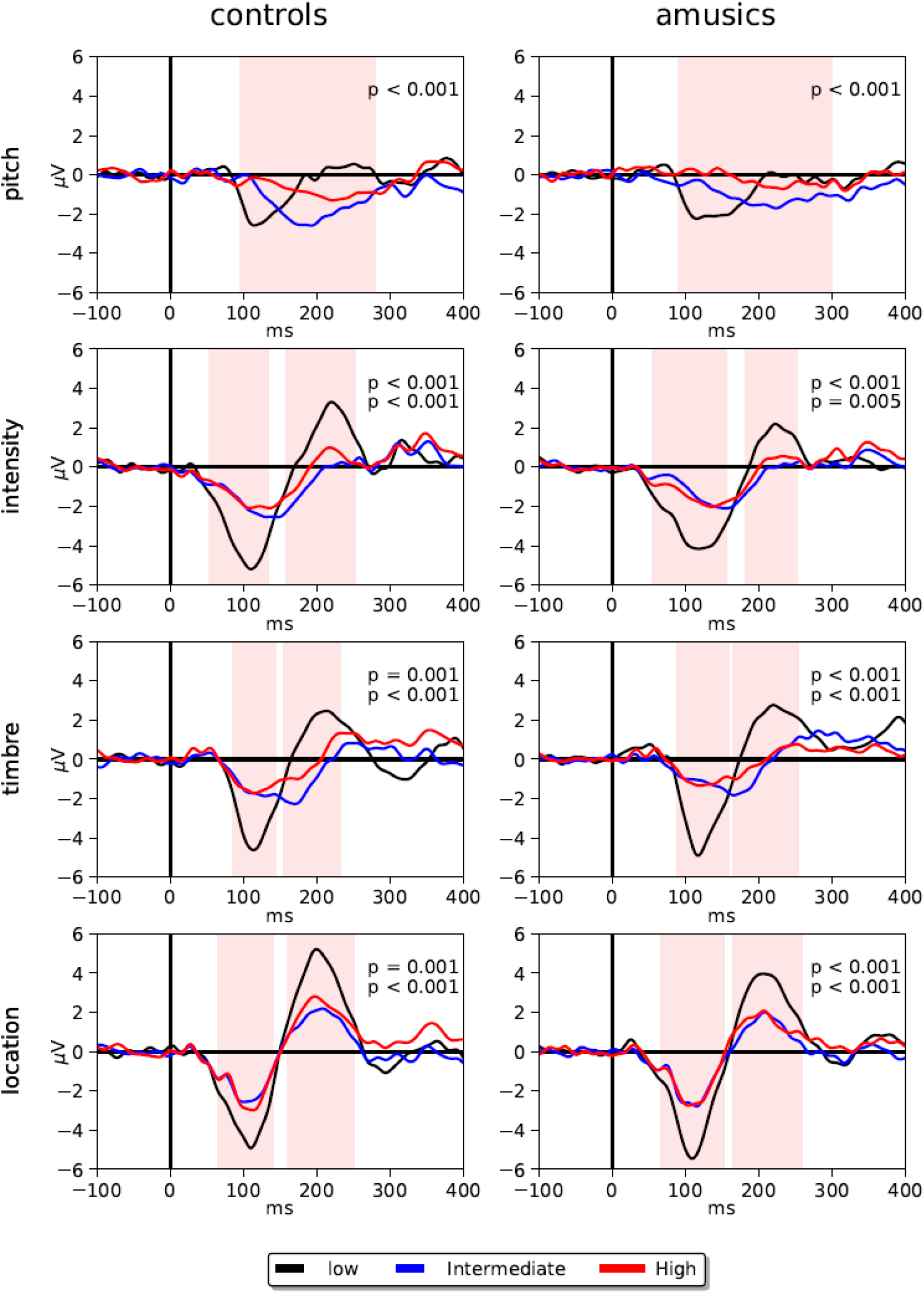
Simple effects of complexity on MMN responses for each group. Shaded areas represent the times when differences between conditions were significant—with a corrected significance threshold of 0.125. The displayed activity corresponds to the average of channels Fz, F1, F2, F3, F4, FCz, FC1, FC2, FC3 and FC4. The given *p*-values correspond to significant clusters.

**Figure 5.**
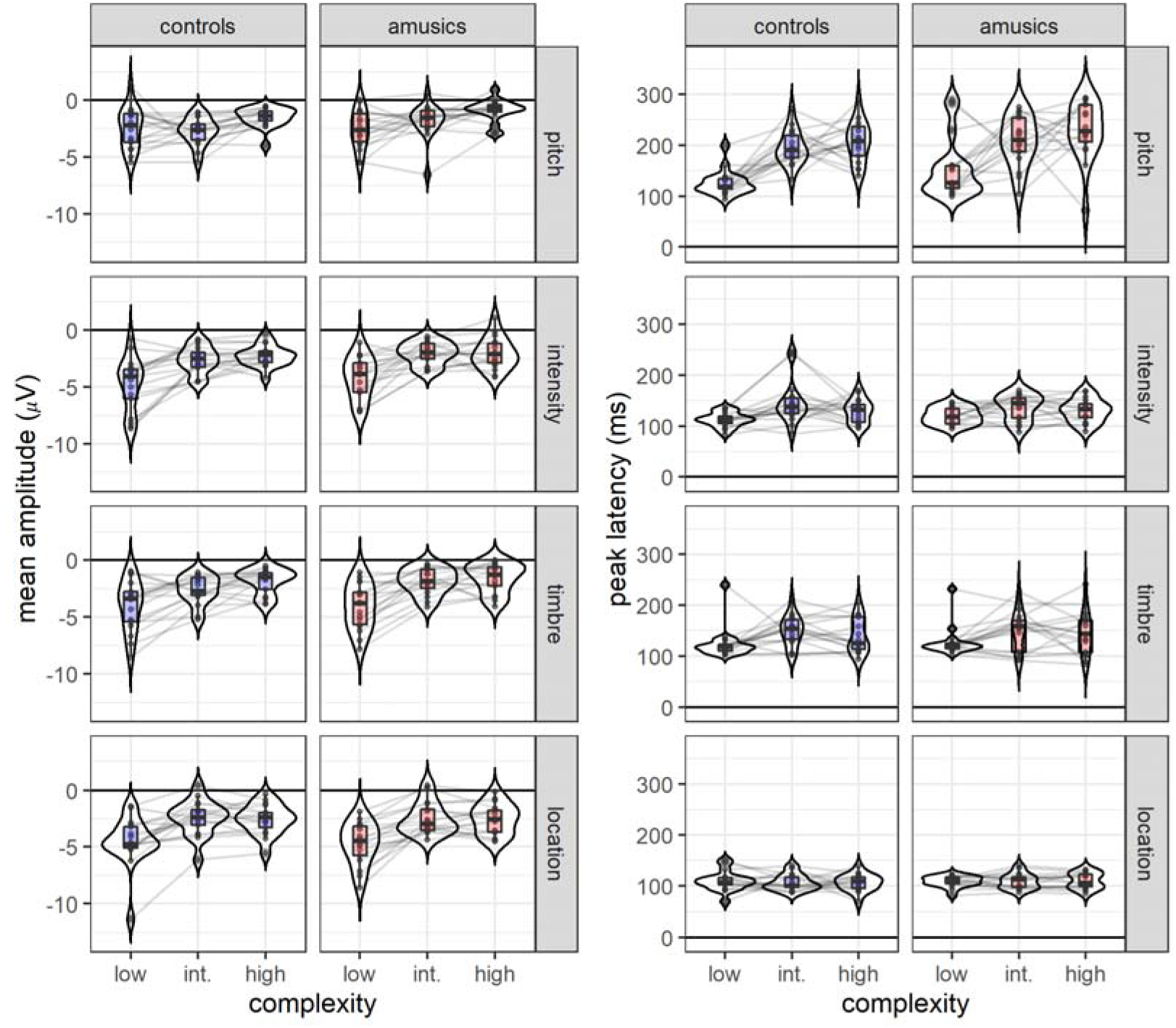
Mean MMN amplitudes (left) and peak latencies (right) as a function of melodic complexity in both groups. Boxes display median and interquartile ranges. Beans depict the estimated densities. Lines connect measurements for individual participants.

Post-hoc, Bonferroni-corrected pairwise contrasts revealed that the complexity effect was mainly driven by significantly larger MMN amplitudes for LC compared with the other two conditions (Table 3). Furthermore, the mean amplitude pairwise contrasts also revealed that pitch deviants followed a different pattern, with significant differences for the comparisons LC-HC and IC-HC, but not LC-IC. Hence the feature-by-complexity interaction.

**Table 3.** Pairwise contrasts of mean amplitudes and peak latencies between complexity conditions. Significant contrasts are highlighted in bold and marked with an asterisk. Standard effect sizes *(d)* are calculated as the difference between conditions divided by the square root of the sum of the residual and the random effects variance.

| feature | contrast | Mean amplitudes |  |  |  |  |  | Peak latencies |  |  |  |  |  |
| --- | --- | --- | --- | --- | --- | --- | --- | --- | --- | --- | --- | --- | --- |
|  |  | est. | CI 2.5% | CI 97.5% | *t* | *p* | *d* | est. | CI 2.5% | CI 97.5% | *t* | *p* | *d* |
| pitch | LC - IC | -0.27 | -0.98 | 0.43 | -0.93 | 1 | -0.18 | -62.15 | -79.68 | -44.61 | -8.52 | **<.001\*** | **-1.92** |
|  | LC - HC | -1.32 | -2.03 | -0.62 | -4.51 | **<.001\*** | **-0.86** | -78.38 | -95.92 | -60.85 | -10.75 | **<.001\*** | **-2.42** |
|  | IC - HC | -1.05 | -1.76 | -0.34 | -3.58 | **<.001\*** | **-0.69** | -16.24 | -33.77 | 1.3 | -2.23 | 0.08 | -0.5 |
| intensity | LC - IC | -2.16 | -2.86 | -1.45 | -7.34 | **<.001\*** | **-1.41** | -24.41 | -41.95 | -6.88 | -3.35 | **<.001\*** | **-0.75** |
|  | LC - HC | -2.32 | -3.03 | -1.61 | -7.9 | **<.001\*** | **-1.51** | -13.15 | -30.68 | 4.39 | -1.8 | 0.22 | -0.41 |
|  | IC - HC | -0.16 | -0.87 | 0.54 | -0.56 | 1 | -0.11 | 11.26 | -6.27 | 28.8 | 1.55 | 0.37 | 0.35 |
| timbre | LC - IC | -1.83 | -2.53 | -1.12 | -6.22 | **<.001\*** | **-1.19** | -24.76 | -42.3 | -7.23 | -3.4 | **<.001\*** | **-0.76** |
|  | LC - HC | -2.34 | -3.05 | -1.64 | -7.98 | **<.001\*** | **-1.53** | -16.38 | -33.92 | 1.15 | -2.25 | 0.08 | -0.51 |
|  | IC - HC | -0.52 | -1.22 | 0.19 | -1.76 | 0.24 | -0.34 | 8.38 | -9.15 | 25.92 | 1.15 | 0.75 | 0.26 |
| location | LC - IC | -2.17 | -2.87 | -1.46 | -7.38 | **<.001\*** | **-1.41** | 1.47 | -16.06 | 19 | 0.2 | 1 | 0.05 |
|  | LC - HC | -2.07 | -2.78 | -1.36 | -7.05 | **<.001\*** | **-1.35** | 1 | -16.53 | 18.53 | 0.14 | 1 | 0.03 |
|  | IC - HC | 0.1 | -0.61 | 0.8 | 0.33 | 1 | 0.06 | -0.47 | -18 | 17.06 | -0.06 | 1 | -0.01 |

For the peak latencies of the MMNs, mixed-effects models revealed effects of complexity (*m*1) and feature (*m*3), as well as a feature-by-complexity interaction (*m*4) (Table 2). Likelihood ratio tests also suggested evidence for a group-by-feature interaction (*m*6), although this result needs to be taken with caution because the *p*-value was close to the significance threshold (*p* = .04) and AIC values indicated *m*4 as the winning model instead. Pairwise contrasts indicated that the complexity-by-feature interaction was mainly driven by larger differences between LC and the other two conditions for pitch deviants than for other deviants (Table 3). Finally, the group-by-feature interaction was driven by an overall longer peak latency of pitch MMNs in amusics compared to controls (Table 4).

**Table 4.**
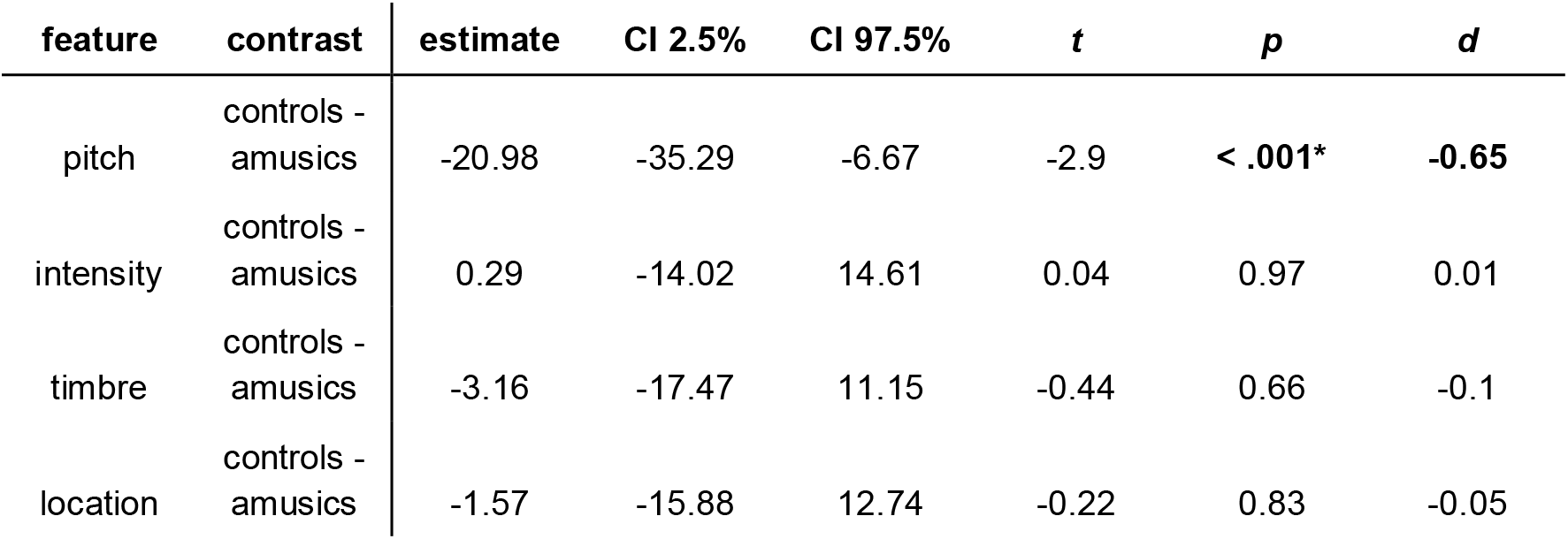
Pairwise contrasts of peak latencies between groups for each feature separately. Significant contrasts are highlighted in bold and marked with an asterisk. Standard effect sizes (*d*) are calculated as the difference between groups divided by the square root of the sum of the residual and the random effects variance.

### 3.3. Familiarity effects

In contrast to complexity, familiarity did not have a significant main effect on MMN amplitudes (Table 5 and see Appendix 4 for cluster level *p*-values). However, in post-hoc analyses of simple effects, significant differences between familiar and unfamiliar pieces emerged for the intensity MMN in the control group (Fig. 6). In amusics, these differences seemed to be present as well, both for pitch and intensity, although they were rendered nonsignificant after multiple comparisons corrections. Interestingly, for amusics, the pitch MMN seemed to disappear in unfamiliar pieces and was significantly smaller than in familiar pieces before correction. Mixed-effects modeling of mean amplitudes revealed a significant main effect of feature (*m*3) (Table 4; Fig 7). Pairwise contrasts indicated that this effect was driven by smaller mean amplitudes of the pitch MMN, and larger mean amplitudes of the location MMN, compared to the other three features (Table 6). Furthermore, AIC values suggested *m*4 as the winning model, which included a feature-by-familiarity interaction. This was supported by a likelihood ratio test with a *p*-value just above threshold (*p* = .067). Pairwise contrasts suggested that this putative effect was related to differences in intensity MMN amplitudes between familiar and unfamiliar stimuli (Table 7). Regarding peak latencies, there was a strong effect of feature, with pitch latencies being longer, and location latencies being shorter than the other three features (Table 6). No significant effect of group or group-related interactions were found in amplitude or latency analyses (Table 5).

**Table 5.** Likelihood ratio tests for mixed-effects models of mean amplitudes and peak latencies for the familiarity conditions. Models were built incrementally by adding a term at the time. Interaction terms are marked with a colon. Comparisons were made between adjacent models. Akaike information criteria (AIC), chi-square statistics (χ^2^) and *p*-values (*p*) are reported. Comparisons with significant differences are highlighted in bold and marked with an asterisk.

| model | added term | null | Mean amplitudes |  |  | Peak latencies |  |  |
| --- | --- | --- | --- | --- | --- | --- | --- | --- |
| | | | AIC | $\chi^2$ | $p$ | AIC | $\chi^2$ | $p$ |
| m0 | Intercept | NA | 874.99 | NA | NA | 2748.63 | NA | NA |
| m1 | familiarity | m0 | 874.28 | 2.71 | 0.1 | 2750.12 | 0.51 | 0.48 |
| m2 | group | m1 | 875.41 | 0.88 | 0.35 | 2751.97 | 0.15 | 0.7 |
| m3 | feature | m2 | 806.63 | 74.78 | **$p < .001^*$** | 2597.67 | 160.3 | **$p < .001^*$** |
| m4 | feature:familiarity | m3 | 805.45 | 7.18 | 0.07 | 2603.42 | 0.25 | 0.97 |
| m5 | group:familiarity | m4 | 806.99 | 0.46 | 0.5 | 2604.34 | 1.08 | 0.3 |
| m6 | feature:group | m5 | 808.77 | 4.23 | 0.24 | 2609.31 | 1.03 | 0.79 |
| m7 | feature:familiarity:group | m6 | 811.28 | 3.49 | 0.32 | 2613.97 | 1.34 | 0.72 |

**Figure 6.**
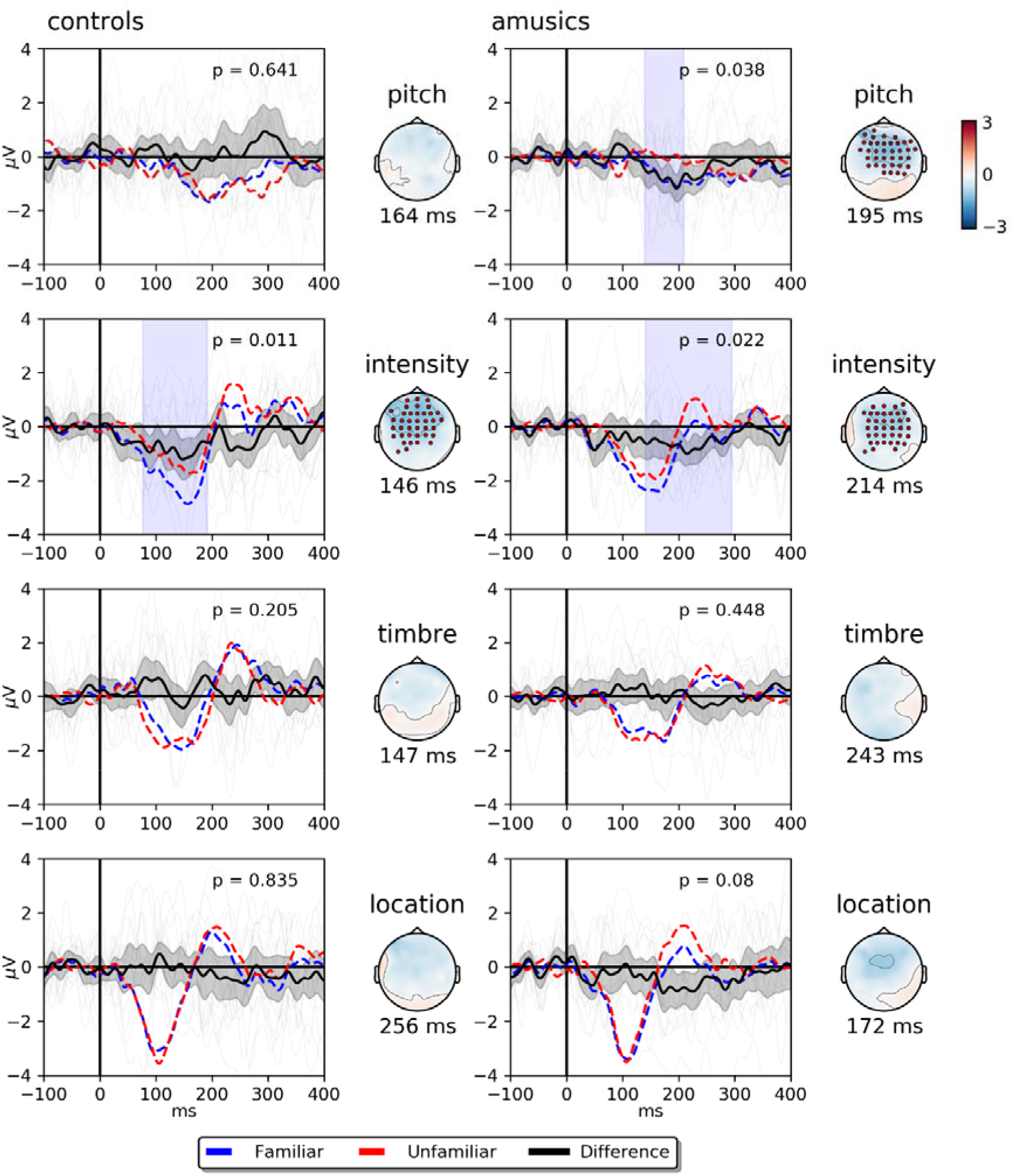
Simple effects of familiarity on MMN responses for each group. Shaded vertical areas represent the times when differences between conditions were significant *before* multiple comparisons correction (our corrected significance threshold is *p* = .0125). Shaded curves depict 95% confidence intervals around the difference between conditions. Grey plot traces correspond to familiarity effects for each participant. Scalp maps show activity at the indicated peak latency of the difference. Red dots represent the channels where differences were significant. The displayed evoked activity corresponds to the average of electrodes Fz, F1, F2, F3, F4, FCz, FC1, FC2, FC3 and FC4. The lowest *p*-value for each feature and group is given.

**Figure 7.**
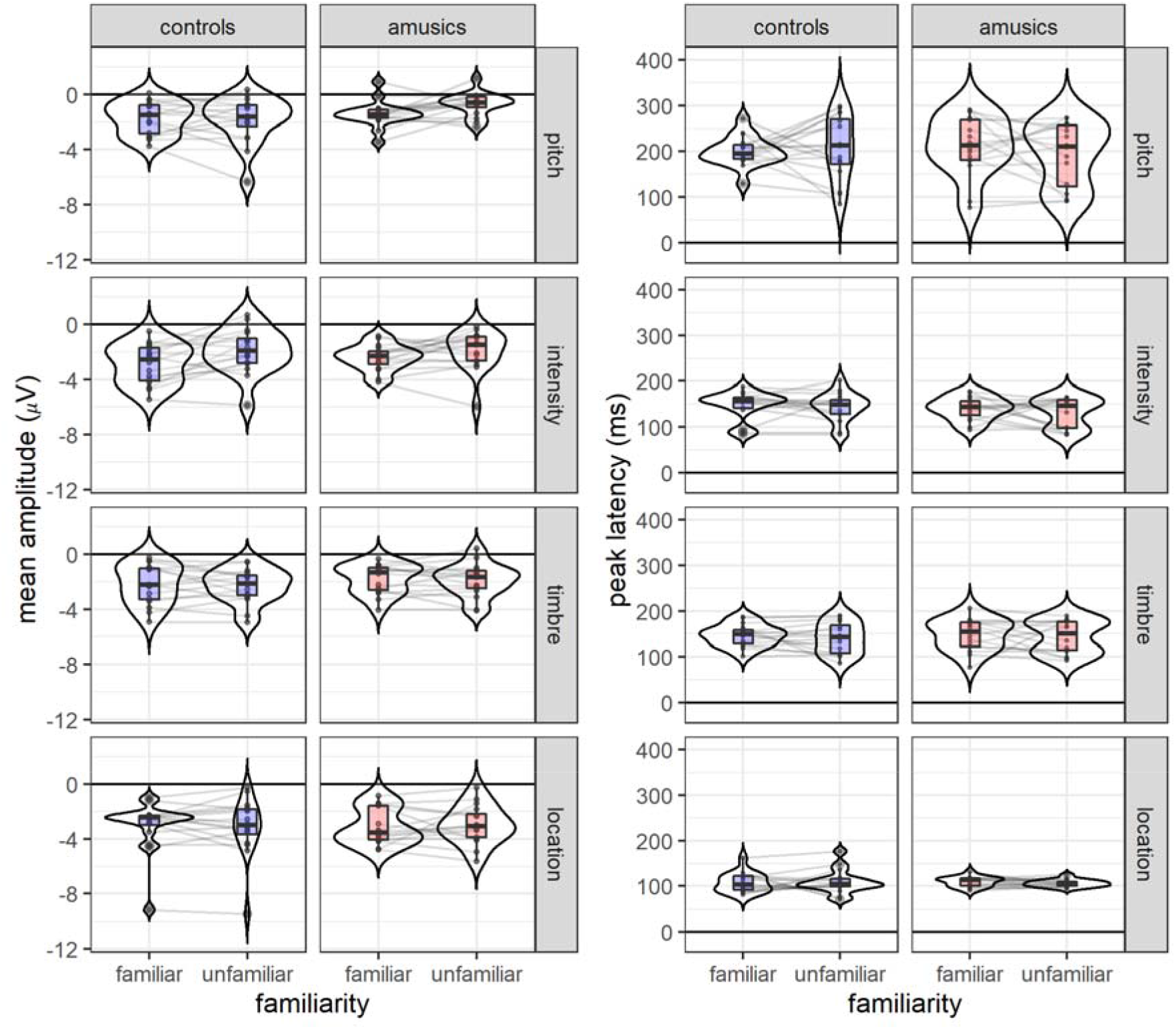
Mean MMN amplitudes (left) and peak latencies (right) as a function of melodic familiarity in both groups. Boxplots display median and interquartile ranges. Beans depict the estimated densities. Lines connect measurements for individual participants.

**Table 6.** Pairwise contrasts of mean amplitudes and peak latencies between features across the familiarity conditions. Comparisons with significant differences are highlighted in bold and marked with an asterisk. Standard effect sizes (*d*) are calculated as the difference between features divided by the square root of the sum of the residual and the random effects variance.

| contrast | Mean amplitudes |  |  |  |  |  | Peak latencies |  |  |  |  |  |
| --- | --- | --- | --- | --- | --- | --- | --- | --- | --- | --- | --- | --- |
| | est. | CI 2.5% | CI 97.5% | $t$ | $p$ | $d$ | est. | CI 2.5% | CI 97.5% | $t$ | $p$ | $d$ |
| pitch - intensity | 0.86 | 0.39 | 1.32 | 4.91 | **<.001\*** | **0.61** | 62.58 | 45.4 | 79.76 | 9.7 | **<.001\*** | **1.62** |
| pitch - timbre | 0.63 | 0.17 | 1.1 | 3.61 | **<.001\*** | **0.45** | 58.81 | 41.63 | 75.99 | 9.12 | **<.001\*** | **1.52** |
| pitch - location | 1.63 | 1.16 | 2.1 | 9.32 | **<.001\*** | **1.15** | 95.39 | 78.21 | 112.57 | 14.78 | **<.001\*** | **2.46** |
| intensity - timbre | -0.23 | -0.69 | 0.24 | -1.3 | 1 | -0.16 | -3.77 | -20.95 | 13.41 | -0.58 | 1 | -0.1 |
| intensity - location | 0.77 | 0.31 | 1.24 | 4.41 | **<.001\*** | **0.55** | 32.81 | 15.63 | 49.99 | 5.09 | **<.001\*** | **0.85** |
| timbre - location | 1 | 0.53 | 1.46 | 5.7 | **<.001\*** | **0.71** | 36.58 | 19.4 | 53.76 | 5.67 | **<.001\*** | **0.94** |

**Table 7.** Pairwise contrasts of mean amplitudes between familiar and unfamiliar melodies, for each feature separately. Comparisons with significant differences are highlighted in bold and marked with an asterisk. Standard effect sizes (*d*) are calculated as the difference between conditions divided by the square root of the sum of the residual and the random effects variance.

| feature | contrast | estimate | CI 2.5% | CI 97.5% | *t* | *p* | *d* |
| --- | --- | --- | --- | --- | --- | --- | --- |
| pitch | familiar - unfamiliar | -0.3 | -0.79 | 0.19 | -1.2 | 0.23 | -0.21 |
| intensity | familiar - unfamiliar | -0.75 | -1.24 | -0.26 | -3.03 | **< .001\*** | **-0.53** |
| timbre | familiar - unfamiliar | 0.11 | -0.38 | 0.6 | 0.45 | 0.65 | 0.08 |
| location | familiar - unfamiliar | -0.03 | -0.52 | 0.46 | -0.12 | 0.9 | -0.02 |

## 4. Discussion

In the present study, we show that the pitch complexity of melodic sequences affects MMN amplitudes and peak latencies for sound feature changes in pitch, intensity, timbre, and location, in both amusic and matched control participants. This demonstrates that, despite their impairment in pitch processing, amusics are sensitive to the contextual uncertainty of pitch sequences. Moreover, we found robust MMN responses for most features and conditions in both groups and no significant interactions between group and complexity or familiarity. This is striking given the predictability range of our stimuli and suggests that preattentive processing of low-level auditory features is far from being fully disrupted in amusics. Nevertheless, we still observed evidence for differences in the processing of pitch deviants between the two groups. In particular, for amusics, the pitch MMN peaked later in the complexity conditions; was not detectable for high complexity and unfamiliar contexts; and was smaller in unfamiliar compared to familiar melodies. In the following, we discuss in detail our findings and their implications, first with regard to predictive uncertainty, and then with regard to how they impact our understanding of congenital amusia.

### 4.1. Effect of pitch complexity on auditory sequence processing

In agreement with previous results (Lumaca et al., 2019; Quiroga-Martinez et al., 2019a, 2019b) we found a reduction in MMN amplitudes with increasing levels of complexity. This is consistent with predictive processing theories and empirical findings suggesting that the strength of prediction error signals is modulated by the uncertainty (or inverse precision) of the context (Clark, 2013; Feldman & Friston, 2010; Garrido et al., 2013; Hohwy, 2012; Hsu et al., 2015; Koelsch et al., 2018; Lumaca et al., 2019; Ross & Hansen, 2016; Sohoglu & Chait, 2016; Southwell & Chait, 2018). In other words, deviant sounds are more salient when auditory sequences are less complex. In Quiroga-Martinez et al. (2019a, 2019b), this effect was observed mainly for pitch-related features, thus pointing to a feature-specificity of the effect. Here, in contrast, the effect was found for all features. This apparent discrepancy can be explained by the introduction of the so-called “optimal” paradigm in the experiment (LC condition), which is based on an oddball sequence and was not present in previous studies.

Note that pairwise differences were found for intensity, timbre and location MMNs between LC and the other two conditions, but not between IC and HC (this lack of difference being in agreement with Quiroga-Martinez et al., 2019a, 2019b). Therefore, a possible explanation is that the use of a single tone in LC stimuli generated, not only higher pitch predictability, but also a stronger sensory representation of the other features. In contrast, in IC and HC conditions, where 31 different tones were employed, the constant change in the acoustic information may have resulted in a somewhat weaker representation of all features. Crucially, IC and HC, but not LC stimuli, are well matched in their acoustic properties and differ only in pitch complexity. This would explain why differences for intensity, timbre and location are found only between LC and the other conditions and why, as suggested by the feature-by-complexity interaction, differences between IC and HC are only present for pitch deviants. This is fully in agreement with previous results (Quiroga-Martinez et al., 2019a, 2019b). Therefore, our findings shed light on complexity effects when oddball sequences are taken into account in the experimental session and suggest that the strength of sensory representations can be disentangled from pitch complexity itself.

The effect of complexity on mean amplitudes was paralleled in peak latencies. In particular for pitch, but also for intensity and timbre, latencies tended to be longer with increasing levels of complexity. This is in agreement with previous research suggesting that the MMN peaks later with decreasing deviance salience due, for example, to deviance magnitude (Näätänen et al., 2007; Sams et al., 1985). Interestingly, location MMNs peaked very early and their latency was not modulated by complexity. This indicates that location information may be processed at very early stages, perhaps even before reaching the auditory cortex, and is less affected by the complexity of the sequences than other features. Furthermore, as in amplitude analyses, significant effects of complexity were found for intensity and timbre only in comparisons between LC and the other conditions, but not between IC and HC. This is in agreement with the hypothesis that sensory representations and pitch complexity effects are dissociable and independent.

Regarding group differences, there was a significant feature-by-group interaction on peak latencies in which the pitch MMN peaked about 20 ms later for amusics than controls, across the complexity conditions. This may be an indication that neural pitch processing is slightly delayed in congenital amusia, in agreement with Albouy et al. (2013). As noted in section 3.3, the evidence for this latency effect is not entirely conclusive and has to be taken with caution. Nevertheless, we believe this finding deserves further exploration as it suggests, for the first time, a pitch-specific difference between amusics and controls in the early and preattentive neural processes indexed by the MMN. Similarly, for high complexity stimuli, there was not a significant pitch MMN response in amusics. This may suggest that amusics are more affected by the complexity of the context. Given that this result goes in the direction of our a priori hypothesis, we believe that it also could be targeted directly in future research.

### 4.2. Effect of melody familiarity on auditory processing

Contrary to our expectations, there were no significant main effects (across groups) of familiarity on MMN amplitudes or latencies for any features. A possible reason for this is that the deviant features that were introduced in the experiment are low-level and depend very little on the abstract representation of a particular melody. After all, the same melody can be played in different instruments (i.e. timbre), with different loudness and reach the ear from different locations. This is particularly interesting in the case of pitch deviants, which consist of mistuned tones, because it suggests that the long-term memory trace of a melody—which is by definition a pitch sequence—is represented independently from the specific tuning with which its pitches are played. Thus, it seems that a given tone in a melody can be recognized even when specific renditions of it are out-of-tune. In this regard, it could be conjectured that familiarity may have an effect only when pitch deviations are equal to or larger than a semitone, i.e. when they actually change the melody and not just the tuning of a given note. Indeed, two previous studies show that this might be the case (Besson et al., 1994; Miranda & Ullman, 2007). These findings are consistent with the idea that listeners perceive music in a categorical way (Goldstone & Hendrickson, 2010; Schulze, 1989; Siegel & Siegel, 1977) and with the feature specificity that underlies multi-feature MMN paradigms (Näätänen et al., 2004; Quiroga-Martinez et al., 2019a; Vuust et al., 2012).

Note, however, that for the intensity MMN there was an indication that amplitudes were larger in the familiar than the unfamiliar condition. In post-hoc, simple-effect comparisons, this pattern was present in both groups, although it was significant only for the control group after correction. The reason for this putative effect is difficult to infer from our experiment, but we may speculate that familiar melodies induce a more pronounced neural tracking of the acoustic envelope that facilitates the detection of intensity changes. Note that pitch and loudness percepts are known to depend on each other (Sek & Moore, 1995) and have been proposed to rely on the same spike-rate code in the auditory cortex (Micheyl et al., 2013). Furthermore, in Quiroga-Martinez et al. (2019a, 2019b), small yet detectable effects of pitch uncertainty on intensity deviants were also reported. Thus, future research on the relationship between pitch uncertainty and intensity MMN responses is warranted.

Our results are surprising when considering previous studies suggesting that MMN responses are enhanced in familiar compared to unfamiliar stimuli (Brattico et al., 2001; Jacobsen et al., 2005; Näätänen et al., 1997). However, in these studies participants were familiar with the type of stimulus itself rather than with how the stimuli unfolded in time. Specifically, it was found that familiarity with the phonemes of a language (Näätänen et al., 1997), a given environmental sound (Jacobsen et al., 2005) or a musical mode (Brattico et al., 2001) enhances mismatch responses. In contrast, here familiarity was defined by how the pitches of a melody follow one another. Therefore, our findings suggest that familiarity with a sound sequence is not sufficient to enhance MMN responses, at least for the deviant features assessed (except intensity) and for typical listeners.

Interestingly, similarly to HC melodies—the most uncertain of the complexity conditions— unfamiliar melodies did not elicit a detectable MMN in amusics. Note that this is in stark contrast with previous experiments that used similarly complex melodies and found intact pitch MMNs in amusics (Zendel et al., 2015, 2009). This discrepancy might be explained by the randomization of the position of the deviant in our experiment, which was not present in the previous studies. In any case, while an interaction between group and familiarity was not significant, this result provides further indications that low-level pitch processing may be impaired in amusics when the context imposes high demands on sensory memory, something that warrants further investigation. This may also be informative regarding previous behavioral research on familiarity effects in congenital amusia, which has revealed preserved effects of familiarity, when assessed with indirect investigation methods targeting implicit processes (Tillmann et al., 2014).

### 4.3. Implications for the understanding of uncertainty and congenital amusia

The results presented here provide insights into two particular issues. First, the fact that MMN responses differed rather little between groups suggests that pre-attentive processing of musical features, and pitch in particular, is far from being fully disrupted in amusia. This is in agreement with the hypothesis that the deficit might be related to abnormal frontotemporal connections that disrupt conscious access to pitch information. Thus, we have shown here for the first time that pre-attentive processing of acoustic changes is sufficiently preserved to allow for an MMN elicitation for different auditory features and for stimuli with varying levels of complexity and familiarity. Interestingly, however, other studies have suggested disrupted encoding of pitch information at the level of auditory cortex (Albouy et al., 2013, 2016, 2019; Tillmann et al., 2016). How can we reconcile the latter with our results and the frontotemporal disconnection hypothesis?

A possible explanation emerges when considering the type of task employed in the different studies. In the “attend” condition of Moreau et al. (2013) and in Peretz et al. (2009) participants had to perform pitch discrimination, which does not necessarily require a thorough encoding of whole pitch sequences. In Albouy et al. (2013), on the other hand, participants had to actively encode a pitch sequence and store it in memory, something that may have required higher top-down input from frontal to temporal areas. Consequently, a hypothesis arising from our work is that disruption in early, low-level auditory processing is mostly apparent in congenital amusia when the task requires active encoding of auditory sequences, such as in short-term memory tasks.

Note however, that we found indications that low-level processing of pitch information may have been disrupted in amusics in the most demanding conditions of the experiment (i.e. high-uncertainty and unfamiliar melodies). This shows that even in a pre-attentive listening task evidence for impairments can be seen. Thus, the lack of conclusive results in this regard (non-significant group effects) may be related to the limited sample size, which is always a challenging aspect due to the low prevalence of congenital amusia—although bear in mind our sample size is large in comparison to most previous EEG studies in congenital amusia. Therefore, we believe that this issue should be explored in further research efforts that overcome sample size limitations, through collaboration between research teams, for example.

The second main insight comes from the fact that listeners with amusia are sensitive to the uncertainty of pitch sequences to the extent that MMN elicitation is disrupted when uncertainty is high. Considering the frontotemporal disconnection hypothesis, this could potentially suggest that processing pitch under high uncertainty conditions requires an intact frontotemporal loop. Note that source localization analyses in Quiroga-Martinez et al. (2019b) indicated differences in anterior superior temporal gyrus between high and low uncertainty pitch MMN responses, thus suggesting this area and its connectivity with inferior frontal regions as targets of future research on congenital amusia.

### 4.4. Limitations

A limitation of our study is that participants listened passively to the stimuli. While this aimed at isolating preattentive responses such as the MMN, further research is needed to assess the effects of complexity and familiarity on responses related to attention and conscious perception, such as the P3, which have been assessed in previous studies (Moreau et al., 2013; Peretz et al., 2005, 2009; Zendel et al. 2015). In this case, active tasks such as deviance detection could be employed. Another limitation is the baseline contamination due to the different phase of rhythm deviants with respect to standard tones. In future experiments, this could be solved by having a longer inter-onset interval, although notice that this would substantially increase the length of the experiment.

### 4.5. Conclusion

The work presented here demonstrates that listeners with congenital amusia are sensitive to the uncertainty of melodies in a similar way to normal listeners. This is reflected in the fact that MMN responses in both groups were reduced by the increasing complexity or reduced familiarity of the melodic context. Furthermore, our results suggest that early preattentive auditory change detection is greatly spared in individuals with amusia, which is the case not only for pitch, but for different auditory features and across different levels of complexity and familiarity. However, we also found evidence for longer pitch MMN latencies and disrupted pitch MMN responses in amusics, especially when context uncertainty was high, something that warrants future research. When taking into account the frontotemporal disconnection hypothesis of amusia, our findings could potentially suggest that processing pitch under high uncertainty conditions requires an intact fronto-temporal loop.

## Open practices

The code and materials employed to conduct the experiment and analyses presented here can be found at: https://doi.org/10.17605/OSF.IO/JSEU8. Due to data protection regulations, data cannot be publicly shared, but can be made privately available upon reasonable request.

## Declaration of interests

None.

## CrediT authorship contribution statement

David R. Quiroga-Martinez: Conceptualization, Methodology, Software, Formal analysis, Data curation, Writing - original draft, Visualization, Investigation. Barbara Tillmann: Conceptualization, Supervision, Writing - original draft. Elvira Brattico: Conceptualization, Supervision, Writing - original draft. Fanny Cholvy - Investigation, Data curation. Lesly Fornoni - Investigation, Data curation. Peter Vuust: Conceptualization, Methodology, Supervision, Writing - original draft. Anne Caclin: Conceptualization, Methodology, Writing - original draft, Supervision.

## Funding information

This work was conducted in the framework of the LabEx CeLyA (‘‘Centre Lyonnais d’Acoustique”, ANR-10-LABX-0060) and of the LabEx Cortex (‘‘Construction, Cognitive Function and Rehabilitation of the Cortex”, ANR-11-LABX-0042) of Université de Lyon, within the program ‘‘Investissements d’avenir” (ANR-16-IDEX-0005) operated by the French National Research Agency (ANR). DQ, EB, and PV are supported by the Danish National Research Foundation (DNRF 117).

---

**Supplementary table 1.** Mean (± SD) scores for each of the subtests in the MBEA.

|  | amusic | control |
| --- | --- | --- |
| MBEA scale | 21.65 (±3.22) | 27.06 (±2.16) |
| MBEA contour | 21.82 (±2.53) | 28.18 (±1.19) |
| MBEA interval | 19.12 (±2.83) | 27.06 (±1.82) |
| MBEA rhythm | 25 (±3.26) | 27.12 (±1.36) |
| MBEA meter | 19.82 (±5.84) | 24.94 (±4.08) |
| MBEA memory | 24.94 (±3.47) | 28.24 (±1.6) |

## Footnotes

1 Note that, although here we consider the Alberti bass as an intermediate condition, in Quiroga-martinez et al. (2019a, 2019b) it was labeled as “low-entropy” condition, because it was the least complex in those experiments.

2 This corresponds to the high-entropy condition in Quiroga-martinez et al. (2019a, 2019b).

## Notes

https://doi.org/10.17605/OSF.IO/JSEU8

### Competing Interest Statement

The authors have declared no competing interest.

### Summary of Updates

New version of manuscript after one round of peer review.

