## Appendix 1 for "Listeners with congenital amusia are sensitive to context uncertainty in melodic sequences"

#### High complexity, familiar and unfamiliar melodies.

### High complexity melodies (major)

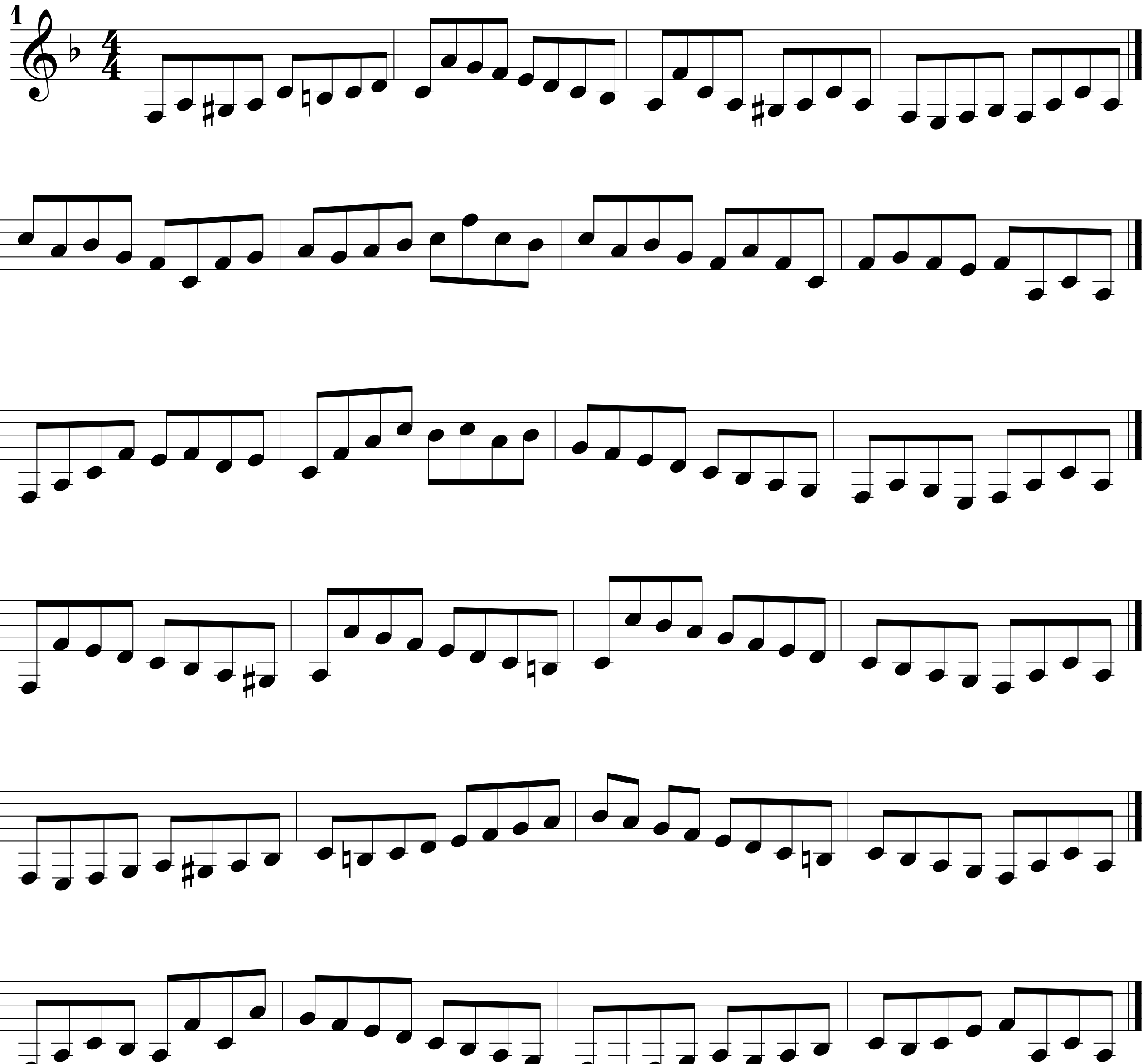


©

### High complexity melodies (minor)

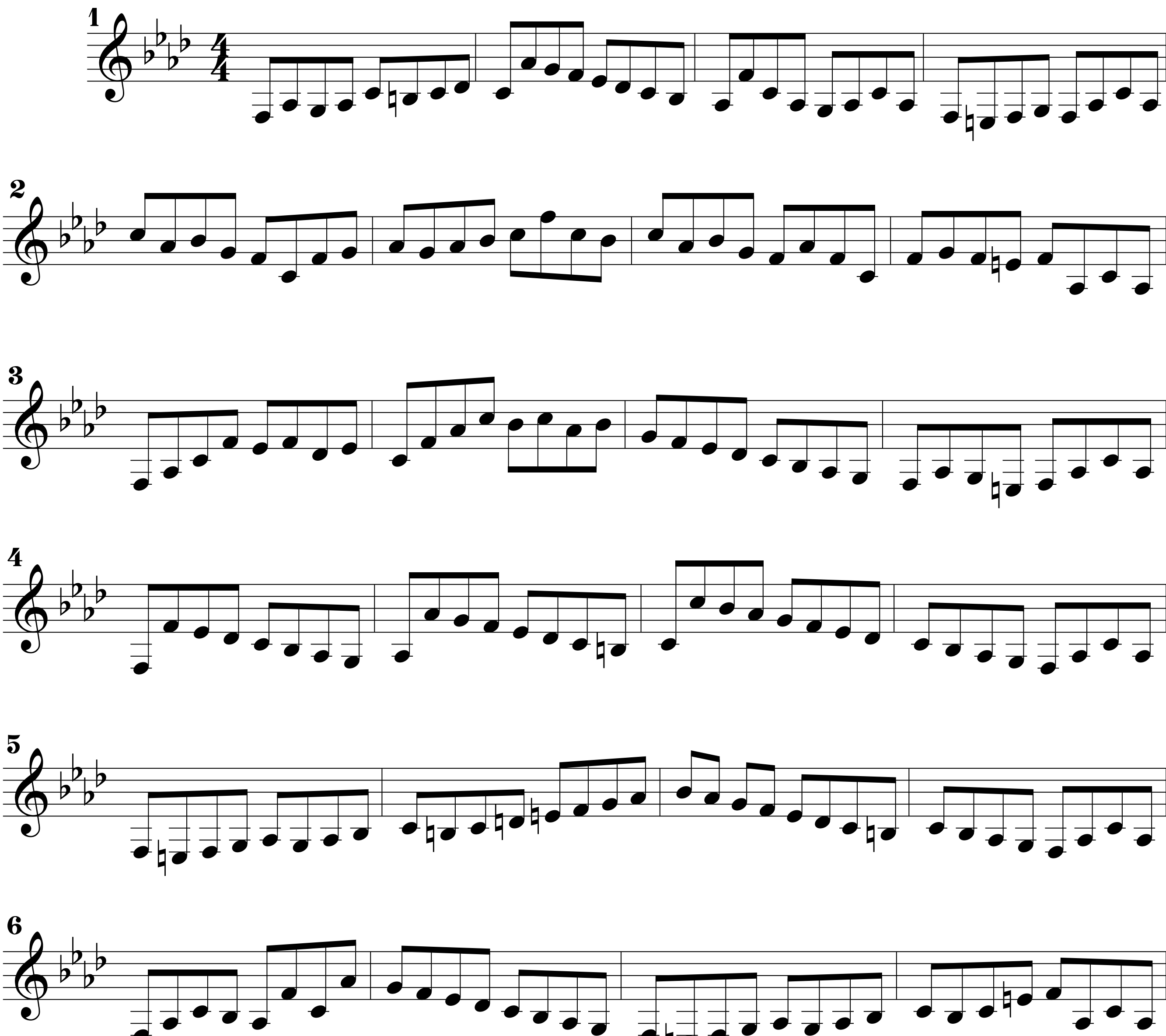


©

### Familiar melodies

Au clair de la lune

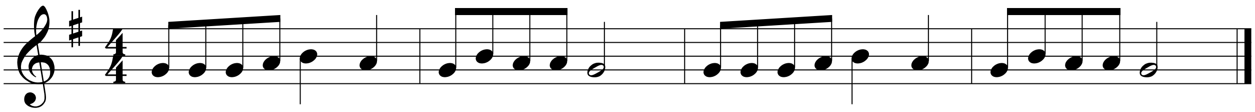

J'ai du bon tabac

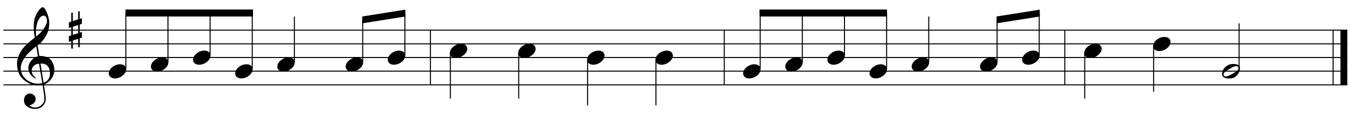

Le pont d'Avignon

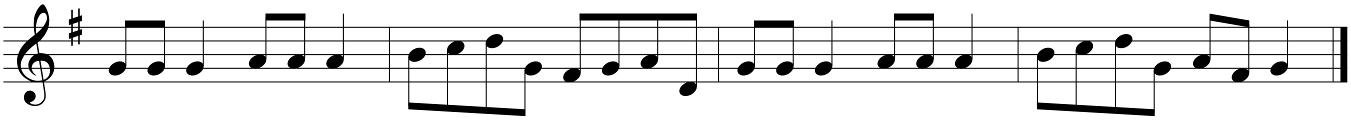

À la claire fontaine

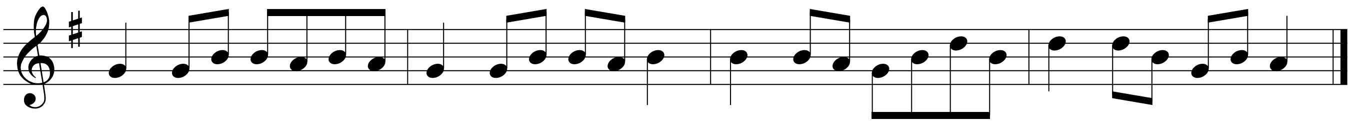

Vive le vent!

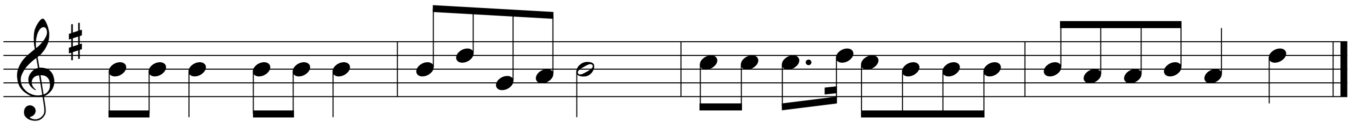

Pomme de reinette et pomme d'api

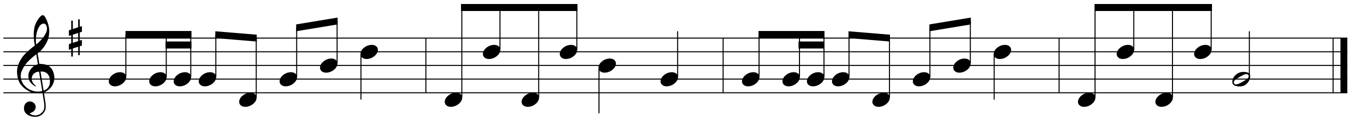

Dodo, l'enfant Do

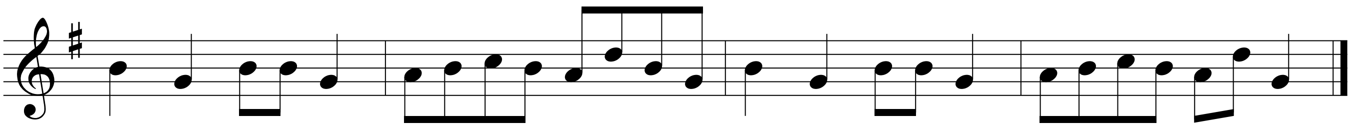

©

### Unfamiliar melodies

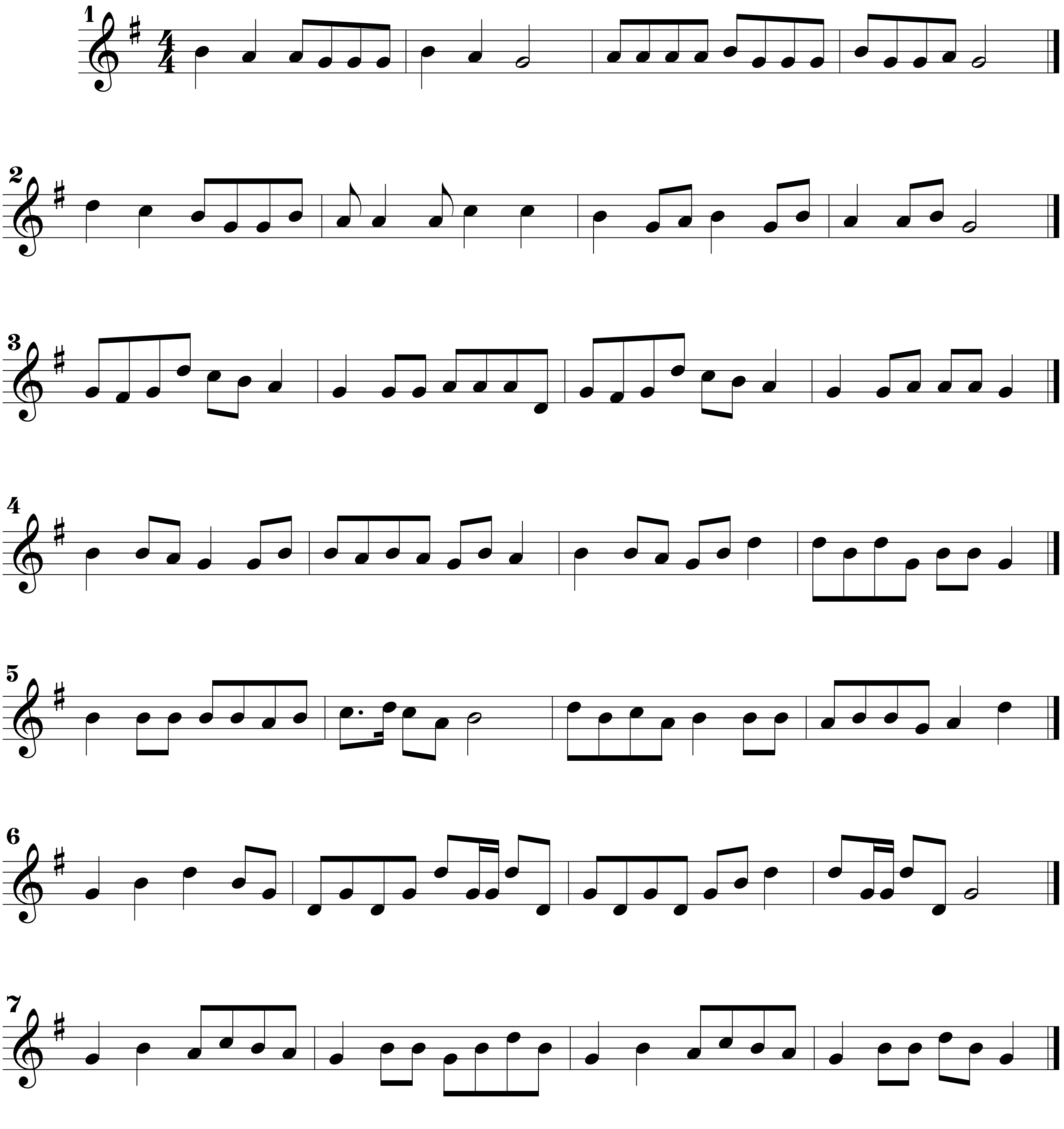


©
