## Appendix 2 for "Listeners with congenital amusia are sensitive to context uncertainty in melodic sequences"

### Appendix 2 − validation of familiarity stimuli

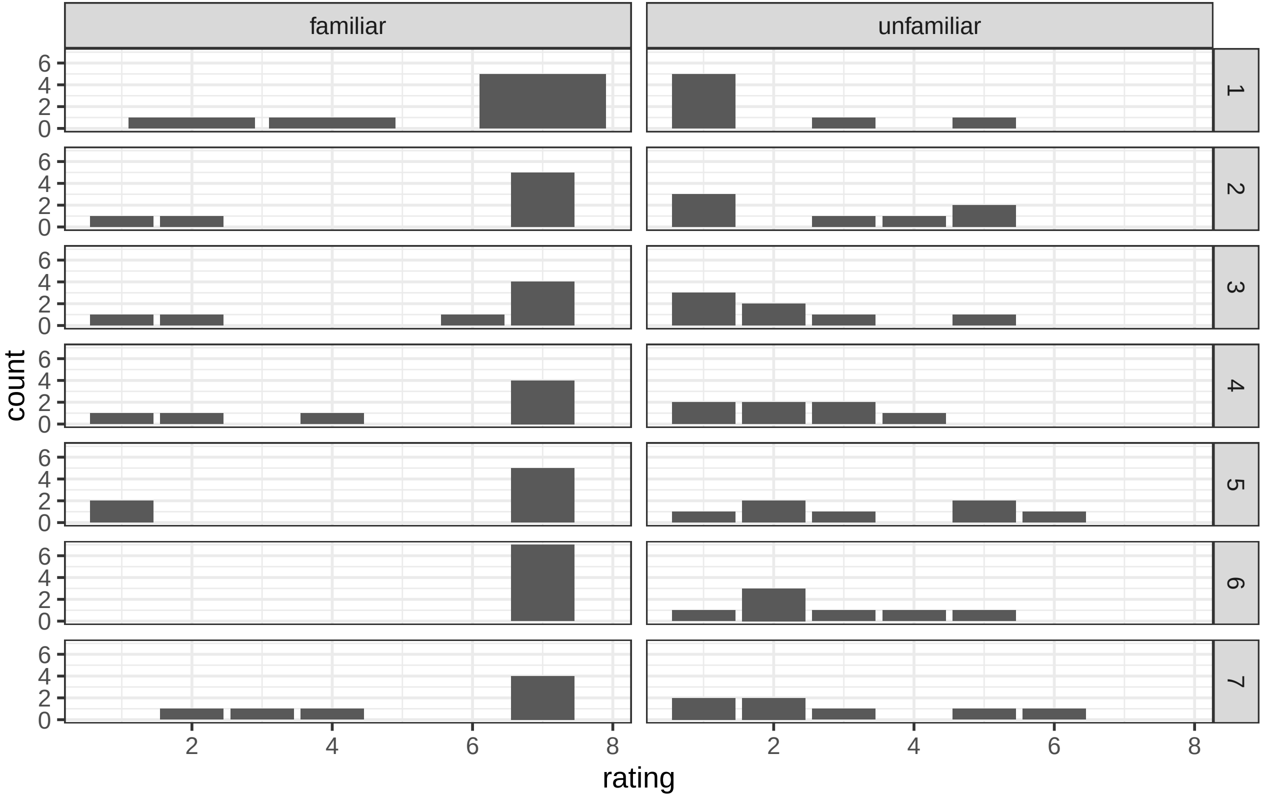


Participants were asked to rate how familiar the melodies were on a scale from 1 to 7. Familiar melodies were rated significantly higher than their unfamiliar scrambled versions (here presented in the same row), as revealed by a cummulative−link mixed−effects model of proportional odds, with melody and subject as random effects ($X^2 = 22.02$, $p < .0001$).
