## Appendix 3 for "Listeners with congenital amusia are sensitive to context uncertainty in melodic sequences"

Standard, deviant and MMN difference waves for rhythm deviations. Note the baseline differences between standard and deviants, which makes them not comparable. This also results in an atypical MMN shape, with large differences before tone onset and after the onset of the next tone. Here the baseline differences have already been ameliorated by baseline correcting the standard ERP with respect to a -160 ms to -60 ms time window. Rhythm MMNs were only detected for intermediate complexity stimuli. Shaded grey areas depict 95% confidence intervals. Shaded blue areas indicate the time points where difference between standard and deviants were significant. Grey traces depict individual MMN waves. Highlighted red dots in the scalp maps indicate the channels where differences were significant. LE, IC, HE = low, intermediate and high complexity. The color range of the scalp map goes from -3 μV (blue) to 3 μV (red).

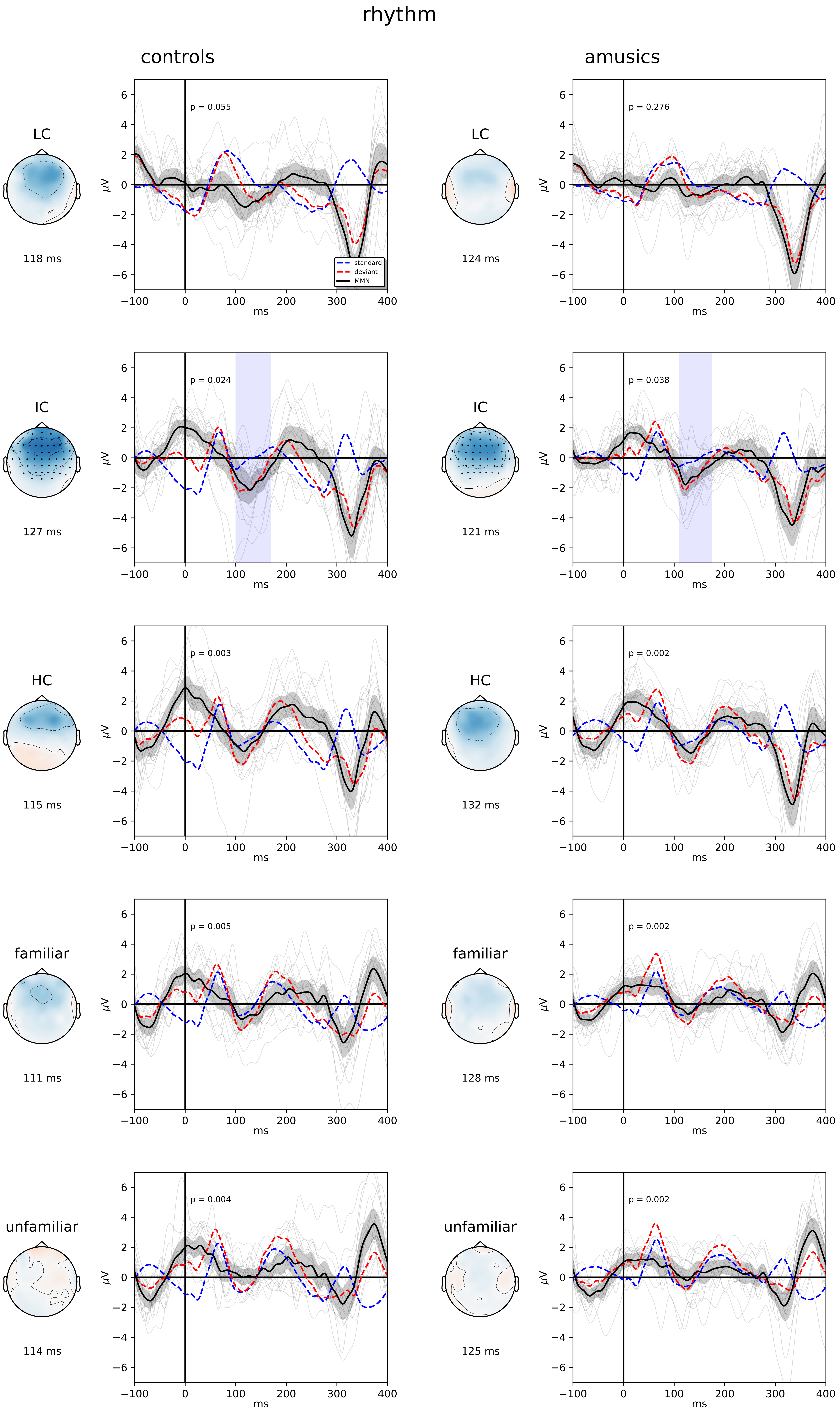
rhythm
controls
amusics
LC
118 ms
p = 0.055
standard
deviant
MMN
LC
124 ms
p = 0.276
IC
127 ms
p = 0.024
IC
121 ms
p = 0.038
HC
115 ms
p = 0.003
HC
132 ms
p = 0.002
familiar
111 ms
p = 0.005
familiar
128 ms
p = 0.002
unfamiliar
114 ms
p = 0.004
unfamiliar
125 ms
p = 0.002
μV
ms
