## Supplementary figures and images for "Listeners with congenital amusia are sensitive to context uncertainty in melodic sequences"

### Appendix 4

#### Appendix 4 - Main effects

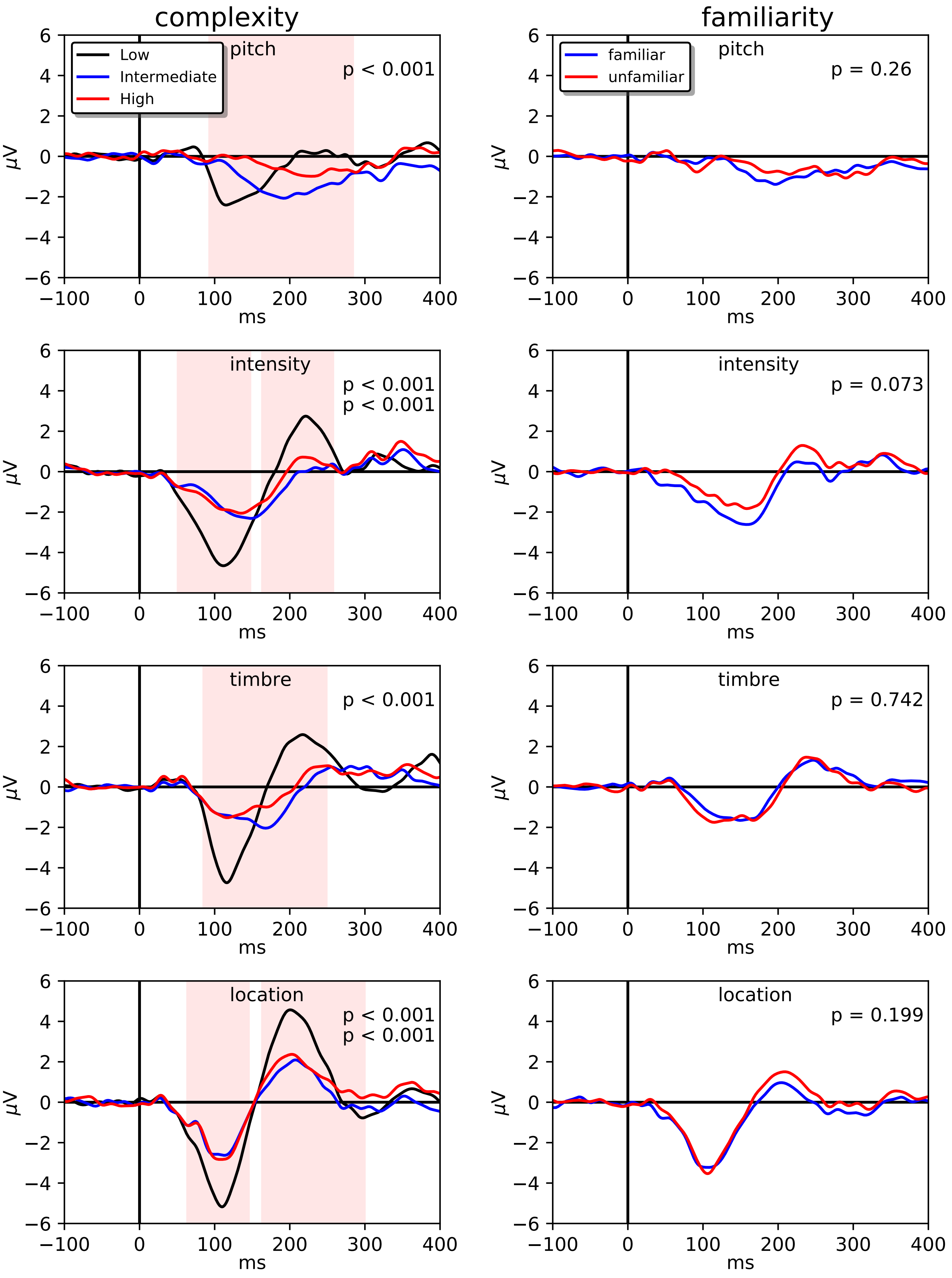
