## Supplemental table 1 for "Listeners with congenital amusia are sensitive to context uncertainty in melodic sequences"

**Supplementary table 1.** Mean (± SD) scores for each of the subtests in the MBEA.

|  | amusic | control |
| --- | --- | --- |
| MBEA scale | 21.65 (±3.22) | 27.06 (±2.16) |
| MBEA contour | 21.82 (±2.53) | 28.18 (±1.19) |
| MBEA interval | 19.12 (±2.83) | 27.06 (±1.82) |
| MBEA rhythm | 25 (±3.26) | 27.12 (±1.36) |
| MBEA meter | 19.82 (±5.84) | 24.94 (±4.08) |
| MBEA memory | 24.94 (±3.47) | 28.24 (±1.6) |
